# Novel engraftment and T cell differentiation of human hematopoietic cells in *Art*^-/-^ *IL2RG*^-/^ SCID pigs

**DOI:** 10.1101/614404

**Authors:** Adeline N Boettcher, Yunsheng Li, Amanda P. Ahrens, Matti Kiupel, Kristen A. Byrne, Crystal L. Loving, A. Giselle Cino-Ozuna, Jayne E. Wiarda, Malavika Adur, Blythe Schultz, Jack J. Swanson, Elizabeth M. Snella, Chak-Sum (Sam) Ho, Sara E. Charley, Zoe E. Kiefer, Joan E. Cunnick, Ellis J. Powell, Giuseppe Dell’Anna, Jackie Jens, Swanand Sathe, Frederick Goldman, Erik R. Westin, Jack C. M. Dekkers, Jason W. Ross, Christopher K. Tuggle

**Affiliations:** Iowa State University, Department of Animal Science, Ames, Iowa; Iowa State University, Laboratory Animal Resources, Ames, Iowa; Michigan State University, Department of Pathobiology and Diagnostic Investigation, College of Veterinary Medicine, East Lansing, Michigan; National Animal Disease Center, Food Safety and Enteric Pathogen Unit, US Department of Agriculture, Agricultural Research Service, Ames, Iowa; Kansas State University, Veterinary Diagnostic Laboratory, Manhattan, Kansas; Immunobiology Graduate Program, College of Veterinary Medicine, Iowa State University, Ames, Iowa; McFarland Clinic, Ames, Iowa; Hope Organ and Tissue Donor Network, Itasca, Illinois; Iowa State University, Veterinary Clinical Sciences, Ames, Iowa; University of Alabama, Department of Pediatrics, Birmingham, Alabama

## Abstract

Pigs with severe combined immunodeficiency (SCID) are an emerging biomedical animal model. Swine are anatomically and physiologically more similar to humans than mice, making them an invaluable tool for preclinical regenerative medicine and cancer research. One essential step in further developing this model is the immunological humanization of SCID pigs. In this work we have generated T^-^ B^-^ NK^-^ SCID pigs through site directed CRISPR/Cas9 mutagenesis of *IL2RG* within a naturally occurring *DCLRE1C* (*Artemis*)^-/-^ genetic background. We confirmed *Art*^-/-^ *IL2RG*^-/*Y*^ pigs lacked T, B, and NK cells in both peripheral blood and lymphoid tissues. Additionally, we and successfully performed a bone marrow transplant on one *Art*^-/-^ *IL2RG*^-/*Y*^ male SCID pig with a bone marrow from a complete swine leukocyte antigen (SLA) matched donor without conditioning to reconstitute porcine T and NK cells. Next, we performed *in utero* injections of cultured human CD34^+^ selected cord blood cells into the fetal *Art*^-/-^ *IL2RG*^-/*Y*^ SCID pigs. At birth, human CD45^+^ CD3ε^+^ cells were detected in peripheral blood of *in utero* injected SCID piglets. Human leukocytes were also detected within the bone marrow, spleen, liver, thymus, and mesenteric lymph nodes of these animals. Taken together, we describe critical steps forwards the development of an immunologically humanized SCID pig model.

**One sentence summary:** We have generated a T^-^ B^-^ NK^-^ SCID pig model through site directed mutagenesis of *IL2RG* in a naturally occurring *Artemis* null background and show successful engraftment of human T and B cells in blood and lymphoid organs after *in utero* injection of human hematopoietic stem cells.

## Introduction

Animals with severe combined immunodeficiency (SCID) are invaluable to biomedical researchers because they are permissive to engraftment of human cells, allowing one to study developmental processes within an *in vivo* environment. In 2012, we discovered the first naturally occurring SCID pigs^1,2^, caused by mutations within the *Artemis* gene, resulting in a T^-^ B^-^ NK^+^ SCID phenotype^3,4^. Since then, pigs with mutations in *RAG1*^5,6^, *RAG2*^7,8^, *IL2RG*^9–11^, and *RAG2*/*IL2RG* ^12^ have also been generated through different mutagenic approaches. Within the past few years, such SCID pigs are now being utilized by cancer^13^, disease model^12^, and stem cell therapy^7^ researchers. Biocontainment facilities^14^, isolators^12^, and Cesarean section^15^ techniques have allowed survival of animals, enabling longer term studies. An important step in further developing the SCID pig model is to immunologically humanize these animals through the introduction of human CD34^+^ hematopoietic stem cells. Similarities between human and porcine immune genes^16^ suggest that human immune development would be supported *in vivo* within the pig^17^. Development of such a model could provide researchers with a larger humanized animal for use in cancer^13,17^, HIV, and vaccine development research.

The first SCID mouse, described in 1983^18^, is capable of being humanized by either injection of human peripheral blood leukocytes^19^ or by implantation of human fetal liver, thymus, and/or lymph node tissue^20^. Reconstitution of human immune cell subsets in SCID mice often requires addition of human cytokine genes, humanization of resident mouse immune genes, or administration of developmental cytokines to the mice ^21–24^. However, limitations of mouse models include differences in size, drug metabolism, and disease pathology compared to humans ^25,26^. Thus, one major goal of the SCID pig community is to create an immunologically humanized SCID pig, which would provide a valuable and unique tool for preclinical research, in a more anatomically and/or physiologically relevant animal model.

The most commonly used strain for humanization is the non-obese diabetic (NOD)-SCID-*IL2RG* (NSG) mouse^27^. The NOD mouse background contains polymorphisms within the SIRPA gene, allowing it to bind to human CD47 to transduce a “don’t eat me” signal in mouse myeloid cells to inhibit phagocytosis^28–30^. We have demonstrated that porcine SIRPA also binds to human CD47 to inhibit phagocytosis of human cells^31^, indicating pigs may be permissive to human xenografts, similar to NOD mice. In addition to the SIRPA polymorphism, NSG mice also have a T^-^ B^-^ NK^-^ cellular phenotype. This cellular phenotype can be generated through mutagenesis of genes required for VDJ recombination (i.e., *Artemis* or *RAG1*/*2*), in addition to *IL2RG*. Previous reports show that mouse NK cells negatively impact human cell engraftment in SCID mice^27^. NK cells in *Art*^-/-^ SCID pigs are functional *in vitro* ^4^, and thus we anticipated swine NK cells could also negatively impact human cell engraftment. To deplete NK cells in our current *Art*^-/-^ SCID pig model, we mutagenized *IL2RG* in an *Art*^-/-^ mutant cell line. The resulting pigs are similar to NSG mice in cellular phenotype and are expected to be similar in SIRPA/CD47 dependent phagocytic tolerance^31^.

Here we describe the generation of *Art*^-/-^ *IL2RG*^-/*Y*^ SCID pigs derived by site-directed CRISPR/Cas9 mutagenesis of *IL2RG* in an *Art*^-/-^ fetal fibroblast cell line. Modified *Art*^-/-^ *IL2RG*^-/*Y*^ embryos, derived from somatic cell nuclear transfer, were implanted in gilts via surgical embryo transfer. Piglets were born at full term and confirmed to have the expected T^-^ B^-^ NK^-^ cellular phenotype based on flow cytometry and immunohistochemical (IHC) analysis of blood and lymphoid organs. We next determined if these double mutant pigs could be humanized via the introduction of human CD34^+^ cord blood stem cells. Gestational day 41 *Art*^-/-^ *IL2RG*^-/*Y*^ fetuses were injected with human CD34^+^ cells within the intraperitoneal space by ultrasound guidance and piglets were delivered via Cesarean section at gestational day 119. We probed for human myeloid, lymphoid, and erythroid cells in peripheral blood and lymphoid organs in piglets of up to 7 days of age. We found evidence of human CD45^+^ cell engraftment in several tissues in the *Art*^-/-^ *IL2RG*^-/-^ pigs. Specifically, we detected CD3ε^+^ T and Pax5^+^ B lymphocytes in blood and lymphoid organs. Taken together, we successfully established the first steps towards the generation of a humanized SCID pig model.

## Results

### Generation of Art^-/-^ IL2RG^-/Y^ SCID pigs by site directed CRISPR/Cas9 mutagenesis of Art^-/-^ fetal fibroblasts

We previously described the discovery of naturally occurring *Art*^-/-^ SCID pigs^3^, which we have been able to raise and breed^14,32^ for research purposes. We started with this genetic background for *IL2RG* site-directed mutagenesis. One goal for producing *Art*^-/-^ *IL2RG*^-/*Y*^ SCID pigs is to generate a breeding colony, such that *IL2RG* knockout piglets can be derived through natural birth rather than cloning procedures. In our breeding protocol, this would require bone marrow transplantation of a SCID boar, so he can be raised to sexual maturity and bred to *Art*^-/+^ carrier females. Mutagenizing our existing *Art*^-/-^ line would facilitate producing SCID pigs with matching SLA (swine leukocyte antigen) to carrier animals within our colony. Therefore, we decided to utilize cloned *Art*^-/-^ fibroblasts for *IL2RG* mutagenesis for somatic cell nuclear transfer (SCNT) to generate pigs with these desired genetics. **Fig. 1** shows a schematic for the process of generating *Art*^-/-^ *IL2RG*^-/*Y*^ piglets.

**Figure 1.**
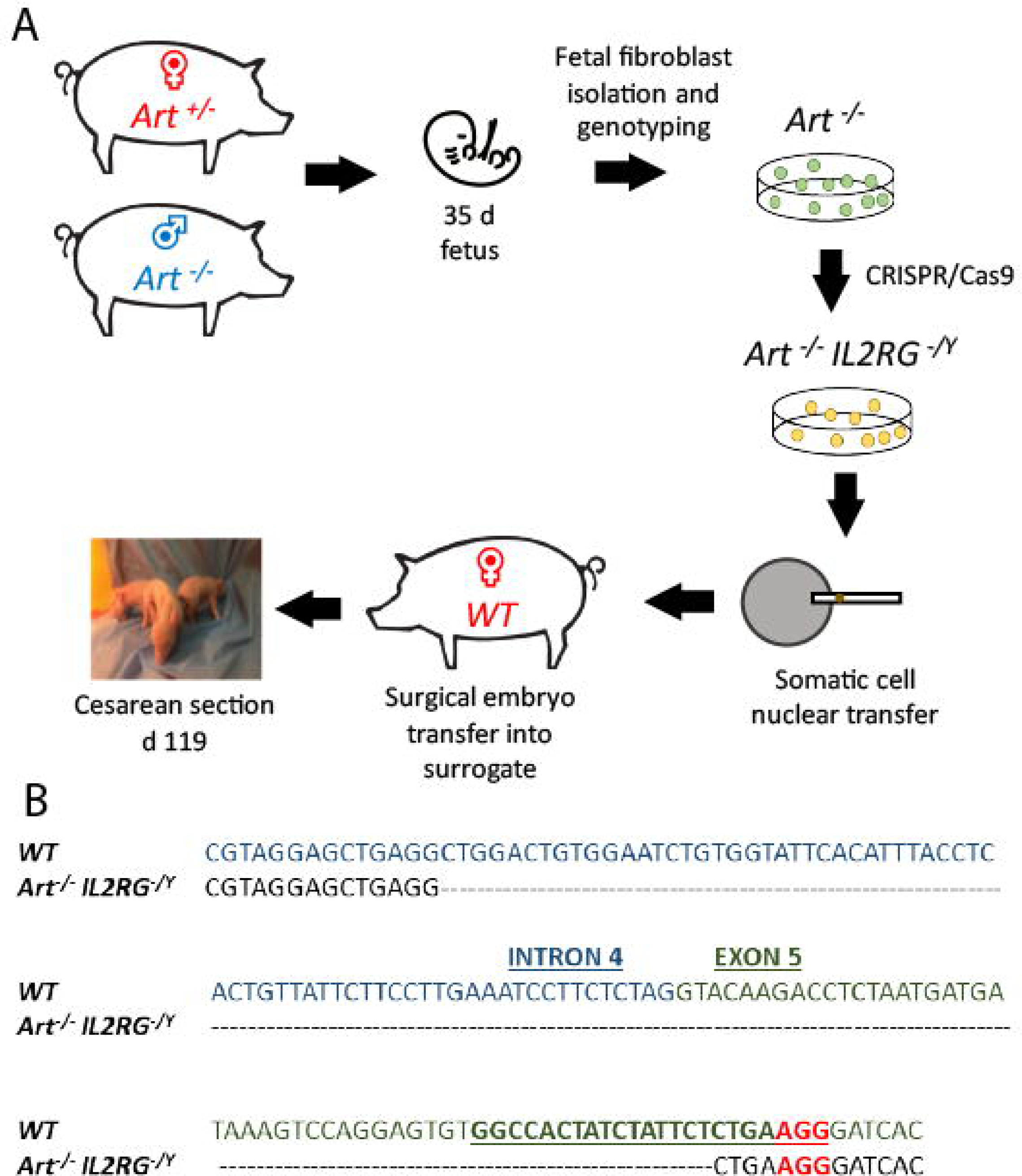
Use of CRISPR/Cas9 system to generate *Art*^*-/-*^ *IL2RG*^*-/Y*^ SCID pigs. **(A)** Semen from an *Art*^-/-^ boar was used to inseminate *Art*^*-/+*^ carrier sows. Two pregnant sows were euthanized at 35 days of gestation and fetal fibroblasts were collected. Collected *Art*^-/-^ fetal fibroblasts were transfected with PX330 plasmid with Cas9 and sgRNA against IL2RG target site, generating *Art*^-/-^*IL2RG*^-/*Y*^ mutant fetal fibroblasts (pFF). Once the mutant pFFs lines were established, somatic cell nuclear transfer was performed, and embryos were transferred into surrogate gilts. Piglets born from these litters were delivered via Cesarean section at gestational day 119 into biocontainment facilities. **(B)** Sequence of IL2RG from wildtype and mutated line. Blue sequence is intron 4, and green sequence is exon 5. Mutation spans through intron 4 and exon 5. Underlined sequence indicates the designed sRNA with the letters in red showing the protospacer adjacent motif (PAM) sequence (NGG).

*Art*^-/-^ pFFs for gene editing were derived from an *Art*^-/-^ male by *Art*^-/+^ female mating (see Materials and Methods for information on *Artemis* genotypes) **(Fig 1A)**. Gestational day 35 fibroblasts were collected and underwent SLA typing and were genotyped for *Art* status. Genotyping revealed that two male and two female cell lines were *Art*^-/-^ mutants, while the other six male and three female cell lines were *Art*^-/+^ carriers (**Supplemental Table 1)**

**Table 1.**
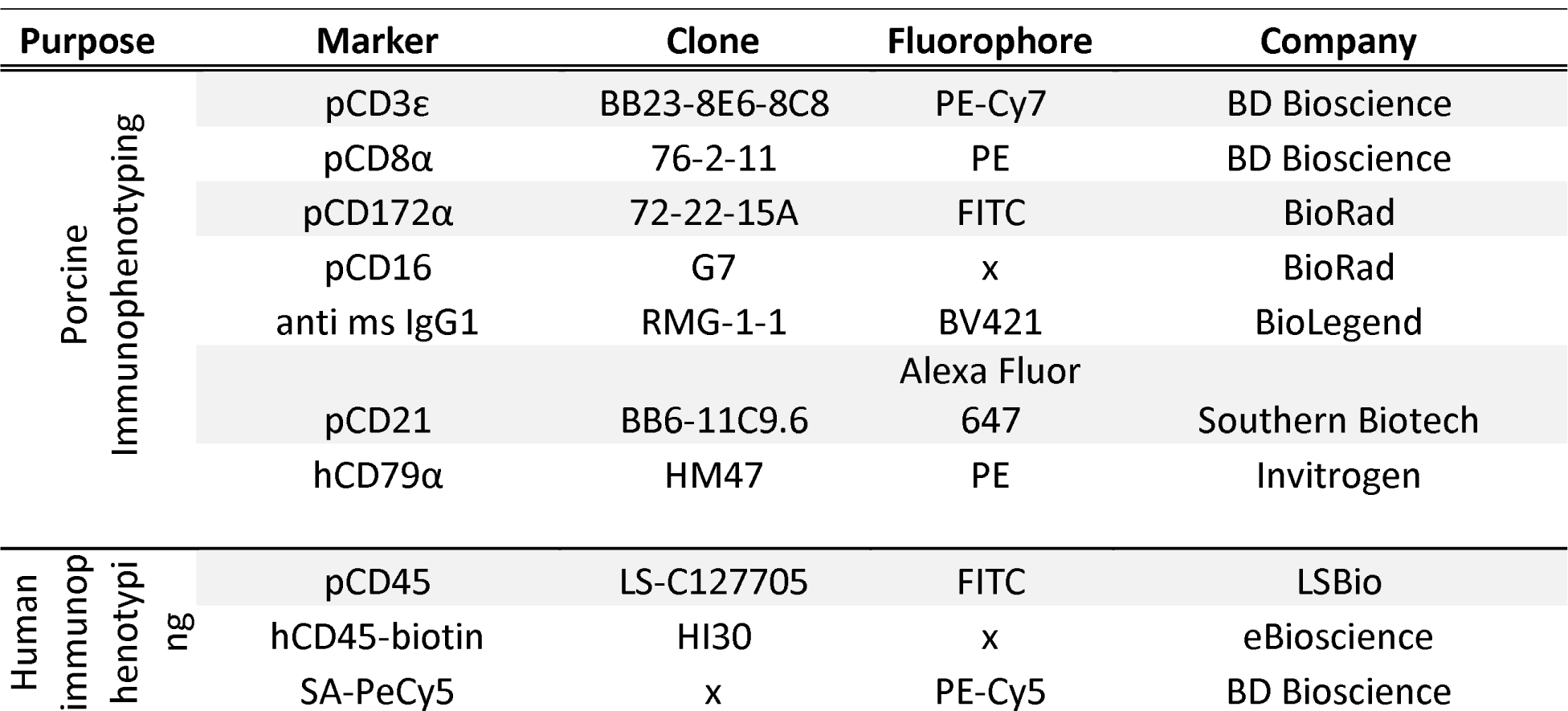
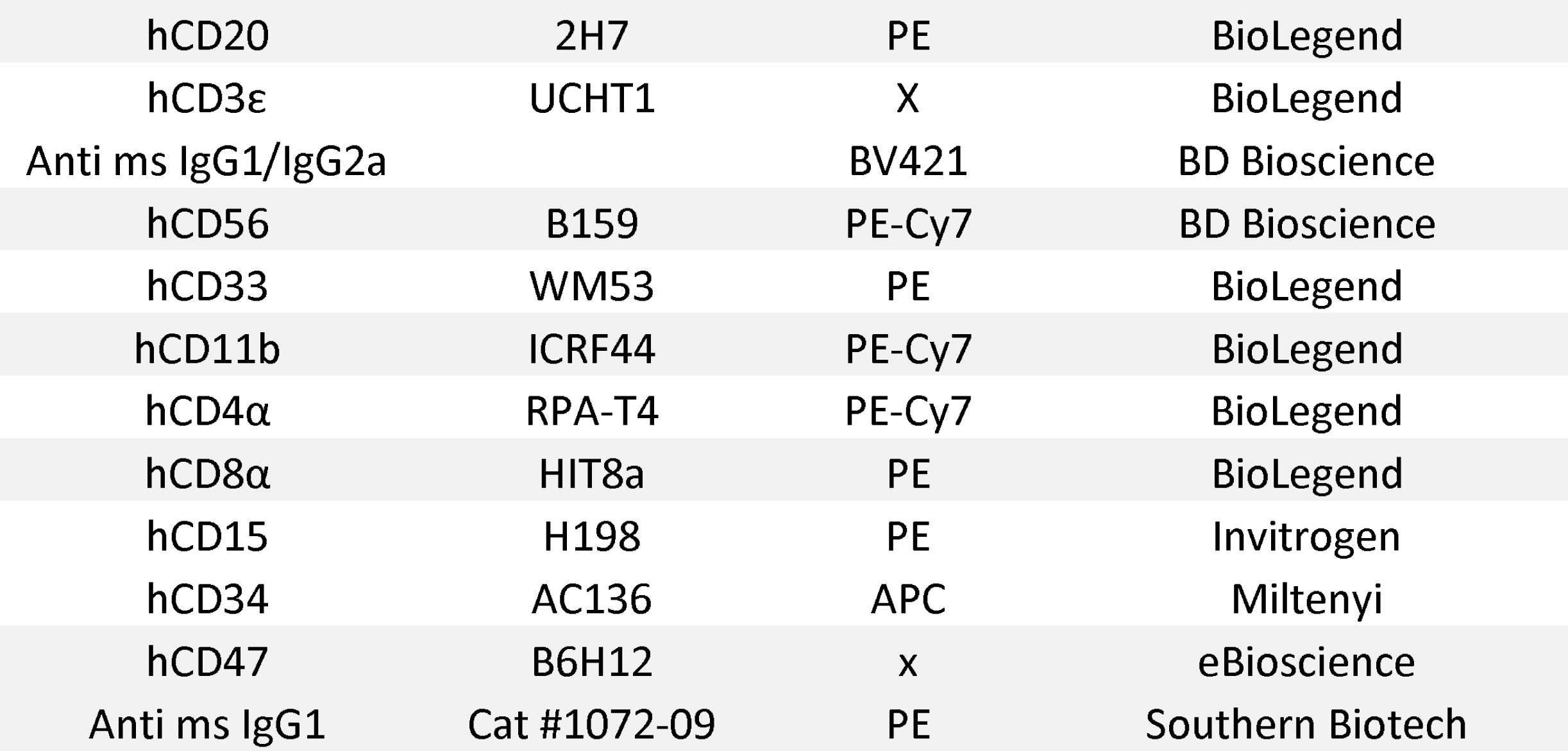
Flow cytometry antibodies used for assessing pig and human cell subsets

| Purpose | Marker | Clone | Fluorophore | Company |
| --- | --- | --- | --- | --- |
| Porcine<br>Immunophenotyping | pCD3ε | BB23-8E6-8C8 | PE-Cy7 | BD Bioscience |
|  | pCD8α | 76-2-11 | PE | BD Bioscience |
|  | pCD172α | 72-22-15A | FITC | BioRad |
|  | pCD16 | G7 | x | BioRad |
|  | anti ms IgG1 | RMG-1-1 | BV421 | BioLegend |
|  | pCD21 | BB6-11C9.6 | Alexa Fluor 647 | Southern Biotech |
|  | hCD79α | HM47 | PE | Invitrogen |
| Human<br>immunop<br>henotypi<br>ng | pCD45 | LS-C127705 | FITC | LSBio |
|  | hCD45-biotin | HI30 | x | eBioscience |
|  | SA-PeCy5 | x | PE-Cy5 | BD Bioscience |

|  |  |  |  |
| --- | --- | --- | --- |
| hCD20 | 2H7 | PE | BioLegend |
| hCD3ε | UCHT1 | X | BioLegend |
| Anti ms IgG1/IgG2a |  | BV421 | BD Bioscience |
| hCD56 | B159 | PE-Cy7 | BD Bioscience |
| hCD33 | WM53 | PE | BioLegend |
| hCD11b | ICRF44 | PE-Cy7 | BioLegend |
| hCD4α | RPA-T4 | PE-Cy7 | BioLegend |
| hCD8α | HIT8a | PE | BioLegend |
| hCD15 | H198 | PE | Invitrogen |
| hCD34 | AC136 | APC | Miltenyi |
| hCD47 | B6H12 | x | eBioscience |
| Anti ms IgG1 | Cat #1072-09 | PE | Southern Biotech |

To mutagenize *IL2RG*, a single guide RNA (sgRNA) was designed to target exon 5 of *IL2RG* (**Fig. 1B).** We selected a male *Art*^-/-^ cell line (7707-FB1) that had a complete SLA-match (haplotype 26.6/68.19a) to a male carrier fibroblast line (7709-FB6) to transfect with a vector to express sgRNA and Cas9 protein **(Supplemental Table 1A, B)**. After transfection, a total of 202 individual clonal colonies from the 7707-FB1 cell line were screened using PCR and Sanger sequencing. Five (2.5 %) clonal colonies were confirmed to be *IL2RG*^-/*Y*^ (hemizygous) **(Supplemental Table 2),** with one cell line (7707-FB1-U23) carrying a 120 bp deletion in intron 4 and exon 5 of the *IL2RG* locus, which was expected to cause a frameshift leading to an premature stop codon **(Fig. 1B)**. A total of 920 *IL2RG*^-/*Y*^ *Art*^-/-^ (7707-FB1-U23) and 512 non-modified (7709-FB6) SCNT derived embryos were transferred surgically into seven recipient gilts. Both cell lines shared the same SLA haplotype of 26.6/68.19a. Among those transferred, four gilts were confirmed pregnant and two carried their piglets to full term and produced five live male piglets via Cesarean section that were reared in biocontainment facilities^14^ (**Supplemental Table 3**). All five piglets were confirmed to have the 120 bp loss in *IL2RG* and the established mutation in exon 10 of *Artemis*^3^ (*Art12* allele information found in Materials and Methods). None of the live born piglets were derived from non-modified, carrier embryos (7709-FB6) that would have been used as a bone marrow donor.

### Art^-/-^ IL2RG^-/Y^ SCID pigs lack T, B, and NK cells in blood and lymphoid organs

Once *Art*^-/-^ *IL2RG*^-/*Y*^ piglets were delivered, blood was collected and analyzed by flow cytometry to confirm the expected T^-^ B^-^ NK^-^ cellular phenotype of these animals. Forward and side scatter plots (FSC/SSC) show the lack of a lymphocyte population in *Art*^-/-^ *IL2RG*^-/*Y*^ pigs compared to *Art*^-/-^ and wild type pigs **(Fig 2A)**. Peripheral blood was stained for pCD172α, pCD8α, and pCD3ε in wildtype, *Art*^-/-^, and *Art*^-/-^ *IL2RG*^-/*Y*^ pigs. Compared to wildtype, T cells (CD172α^-^ CD3ε^+^) and NK cells (CD172α^-^ CD3ε^-^ CD8α^+^) were not in the circulation of *Art*^-/-^ *IL2RG*^-/*Y*^ piglets **(Fig. 2B)**. We additionally stained for pCD21 to confirm there were no B cells in the circulation of *Art*^-/-^ *IL2RG*^-/*Y*^ pigs **(Supplemental Fig. 1)**. Furthermore, *Art*^-/-^ *IL2RG*^-/*Y*^ pigs had atrophic and smaller thymus, spleen, and lymphoid tissue within the intestines compared to wildtype pigs, (**Supplemental Fig. 2)**.

**Figure 2.**
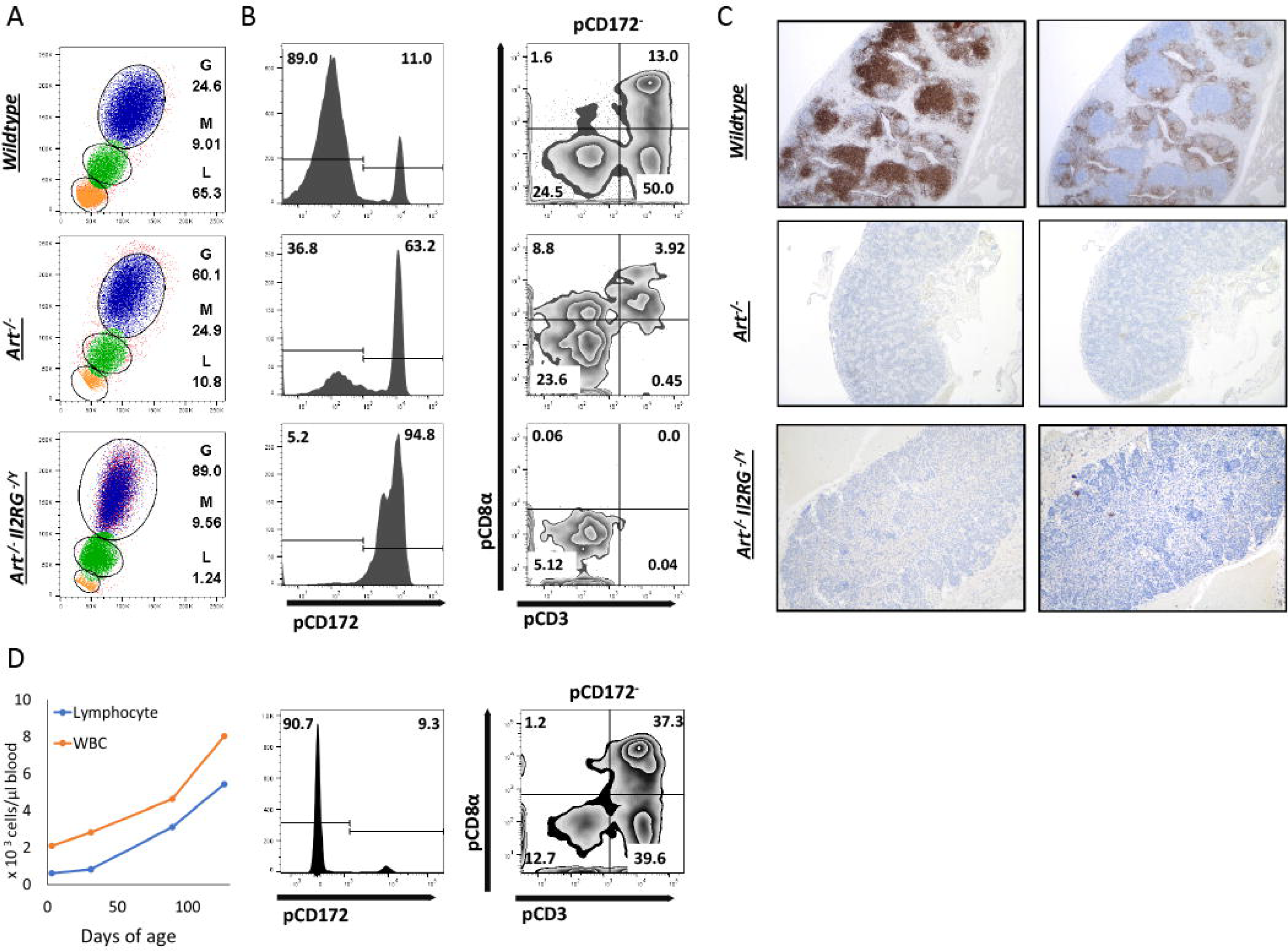
*Art*^-/-^ *IL2RG*^-/*Y*^ SCID pigs lack T, B, and NK cells in peripheral blood and lymphoid organs and T cells can be reconstituted after pig to pig bone marrow transplantation. **(A)** Flow cytometric analysis of major leukocyte populations in wildtype, *Art*^-/-^, and *Art*^-/-^ *IL2RG*^-/*Y*^ pigs. Percentages for granulocytes (G), monocytes (M), and lymphocytes (L) are shown for each type of pig. **(B)** Mononuclear populations were gated from each sample and assessed for presence of monocytes (CD172α^+^ CD3ε^-^ CD8α^-^), T cells (CD172α^-^ CD3ε^+^ CD8α^-/+^), and NK cells (CD172^-^CD3ε^-^ CD8α^+^). CD172α expression was assessed first, then CD172α^-^ populations were assessed for expression of CD3ε and CD8α. *Art*^-/-^ *IL2RG* ^-/*Y*^ pigs lack T and NK cells. **(C)** Lymph nodes were collected and assessed for T cells (CD3ε) and B cells (CD79α). **(D)** One *Art*^-/-^ *IL2RG*^-/*Y*^ SCID piglet underwent a bone marrow transplant at 5 days of age. At 4 months post transplantation, MNC were stained for CD172α, CD3ε, and CD8α to assess for donor T and NK cells (left). Complete blood counts were also performed throughout a four-month period (right). White blood cell and lymphocyte concentrations increased at every collection point.

We then confirmed that T and B cells were absent from lymphoid tissues. Lymph nodes were collected from wild type (18d), *Art*^-/-^ (0d), and *Art*^-/-^ *IL2RG*^-/*Y*^ (18d) pigs for IHC analysis. Lymph node tissue was stained for CD3ε and CD79α to assess for the presence of T and B cells, respectively. Both *Art*^-/-^ and *Art*^-/-^ *IL2RG*^-/*Y*^ pigs lacked normal T and B cells in lymph nodes **(Figure 2C).**

### T and NK cell reconstitution in an Art^-/-^ IL2RG^-/Y^ SCID pig after pig bone marrow transplantation

An additional goal for generation of *Art*^-/-^ *IL2RG*^-/*Y*^ SCID pigs was to establish a male breeding population to maintain this line. Carrier *Art*^-/+^ females bred with an *Art*^-/-^ *IL2RG*^-/*Y*^ SCID boar would generate litters with a mix of males and females with different *Art* and *IL2RG* genotypes for future studies and breeding. The original intent of performing embryo transfers with non-modified carrier embryos was for them to provide a source of bone marrow for the *Art*^-/-^ *IL2RG*^-/*Y*^ pigs. In our two full-term pregnancies, one gilt was transferred with carrier and *Art*^-/-^ *IL2RG*^-/*Y*^ embryos (7707-FB1-U23 and 7709-FB6), while the other was transferred with only *Art*^-/-^ *IL2RG*^-/*Y*^ embryos. From both litters, only Art^-/-^ *IL2RG*^-/*Y*^ piglets were born, thus requiring an alternative source of bone marrow for a BMT.

We therefore performed a BMT on one *Art*^-/-^ *IL2RG*^-/*Y*^ male piglet using complete SLA matched (26.6/68.19a) bone marrow from a 4-year-old sow that was raised on a conventional farm. A total of 2.27 x 10^8^ million unfractionated bone marrow cells from the ribs and sternum were collected and administered to one *Art*^-/-^ *IL2RG*^-/*Y*^ SCID pig. The piglet was not conditioned prior to the BMT, based on our previous success with porcine T and B cell reconstitution without conditioning in *Art*^-/-^ SCID pigs^32^.

Peripheral blood was collected approximately once a month after the BMT and analyzed by either a complete blood count (CBC) or by flow cytometric analysis (FACS). CBC analysis showed that white blood cells and lymphocytes increased monthly after the transplant, to near normal levels by 4 months post BMT **(Fig. 2D)**. FACS analysis using antibodies against pCD3ε, pCD8α, and pCD172α revealed circulating T and NK cells at 4 months of age **(Fig. 2D).** Additionally, we used FACS analysis to probe peripheral blood for CD21 and CD79α expressing cells, and we observed very few circulating B cells 6 months post BMT (**Supplemental Fig. 3**). Of note, B cell reconstitution is variable in human SCIDs with mutations in *Artemis* post BMT^33–35^, which may be a function of conditioning regimens. Importantly, the *Art*^-/-^ *IL2RG*^-/*Y*^ boar post BMT has been maintained in the biocontainment bubble since birth and is sexually mature and healthy as of 11 months of age.

**Figure 3.**
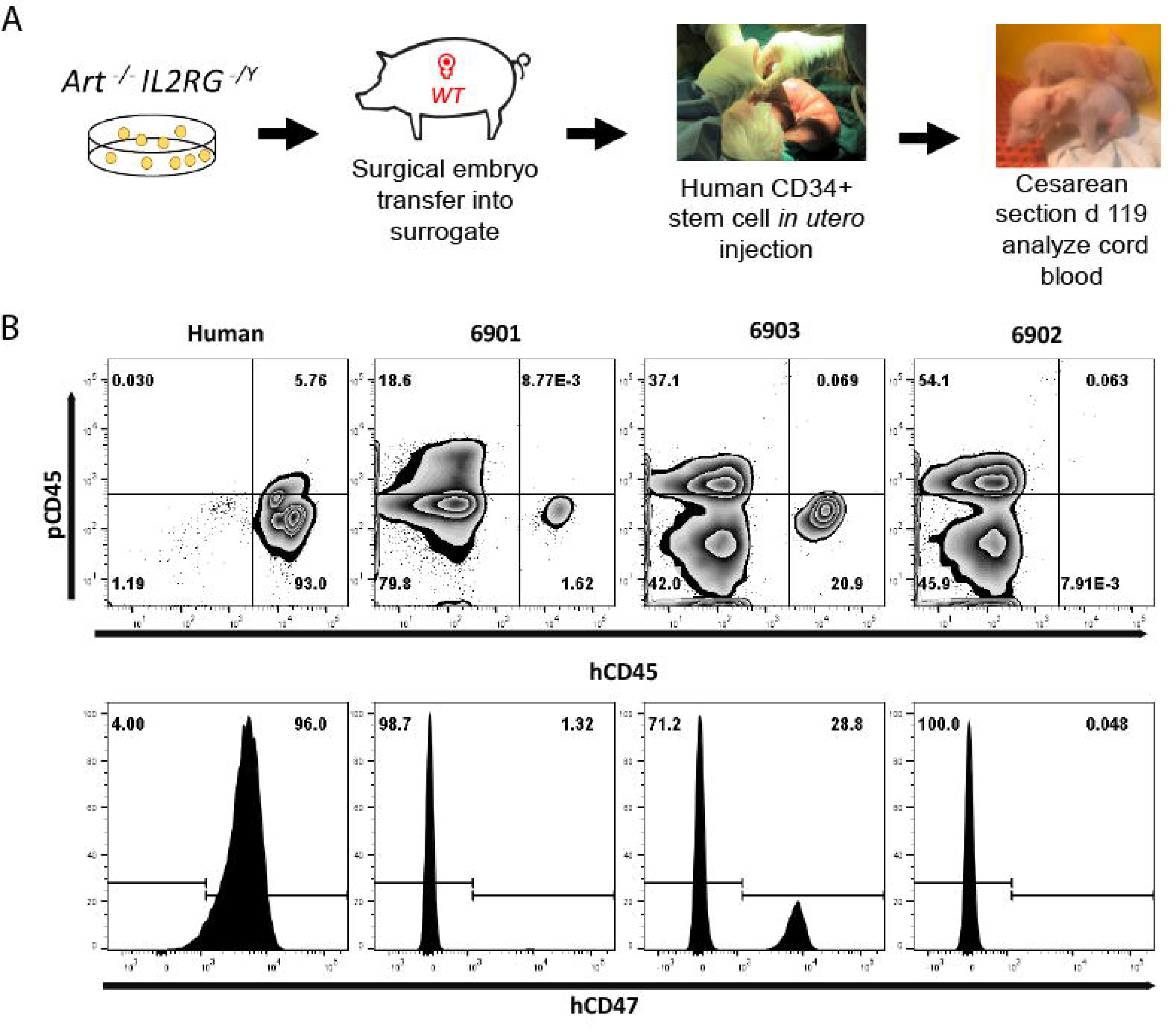
Human leukocytes in *Art*^-/-^ *IL2RG*^-/*Y*^ SCID pig cord blood. **(A)** Somatic cell nuclear transfer derived *Art*^-/-^ *IL2RG*^-/*Y*^ embryos were surgically transferred into a surrogate gilt. At gestational day 41, the pregnant gilt underwent a laparotomy procedure to expose the uterus and ultrasound guidance was utilized to inject human CD34^+^ stem cells into the intraperitoneal space of fetal piglets. After surgery, piglets were delivered via Cesarean section at gestational day 119 into biocontainment facilities. **(B)** Blood was collected from a human and three neonatal (6901, 6902, and 6903) SCID pigs and stained for human and pig CD45, as well as human CD47. All cells in circulation were gated from SCID pigs. Pigs 6901 and 6903 both had human cells in cord blood, as shown by presence of hCD45^+^ and hCD47^+^ cells.

### Circulating human T cells in neonatal Art^-/-^ IL2RG^-/Y^ piglets after in utero injection of human hematopoietic stem cells

Once we confirmed the cellular phenotype of *Art*^-/-^ *IL2RG*^-/*Y*^ pigs, we investigated whether these pigs were capable of engrafting human CD34^+^ hematopoietic stem cells. We had previously attempted intravenous or intraosseous injection of human CD34^+^ cells into single mutant *Art*^-/-^ piglets (within one week of age), with various cell doses and busulfan conditioning **(Supplemental Table 4)** and we did not detect any evidence of engraftment in peripheral blood or lymphoid organs fifteen weeks post-transplant, there was no evidence of engraftment within blood or lymphoid organs. As an alternative approach, *in utero* injection of human stem cells into the intraperitoneal space of *Art*^-/-^ *IL2RG*^-/*Y*^ SCID pig fetuses was performed

During mid-gestation (30-45 days), the fetal liver is the major site for hematopoiesis for many species^36–39^, including swine^40^. Previous reports show that injection of human CD34^+^ cells into the intraperitoneal space of fetal piglets and sheep leads to differentiation and engraftment of human immune cells^41–44^. Engraftment of human cells into pig and sheep fetuses is facilitated by the fact that the cellular environment is immunologically privileged early in gestation. However, such injections have not been reported in SCID pigs. Approaches to inject fetal piglets require laparotomy procedures to expose the uterus to visualize fetuses via ultrasound imaging. Thus, laparotomy surgery with ultrasound guidance was used to inject human CD34^+^ cells into the intraperitoneal space of *Art*^-/-^ *IL2RG*^-/*Y*^ fetuses ^45^ **(Fig. 3A)**. Positively selected human CD34^+^ cells isolated from cord blood were cultured with thrombopoietin (TPO), FLT-3L, and stem cell factor (SCF) for the seven days prior to injection to enhance expansion for the appropriate cell dose delivery. After culturing, cells were either CD45^+^CD34^+^ (23-28%) or CD45^+^ CD34^-^ (71-76%) **(Supplemental Figure 4).**

**Figure 4.**
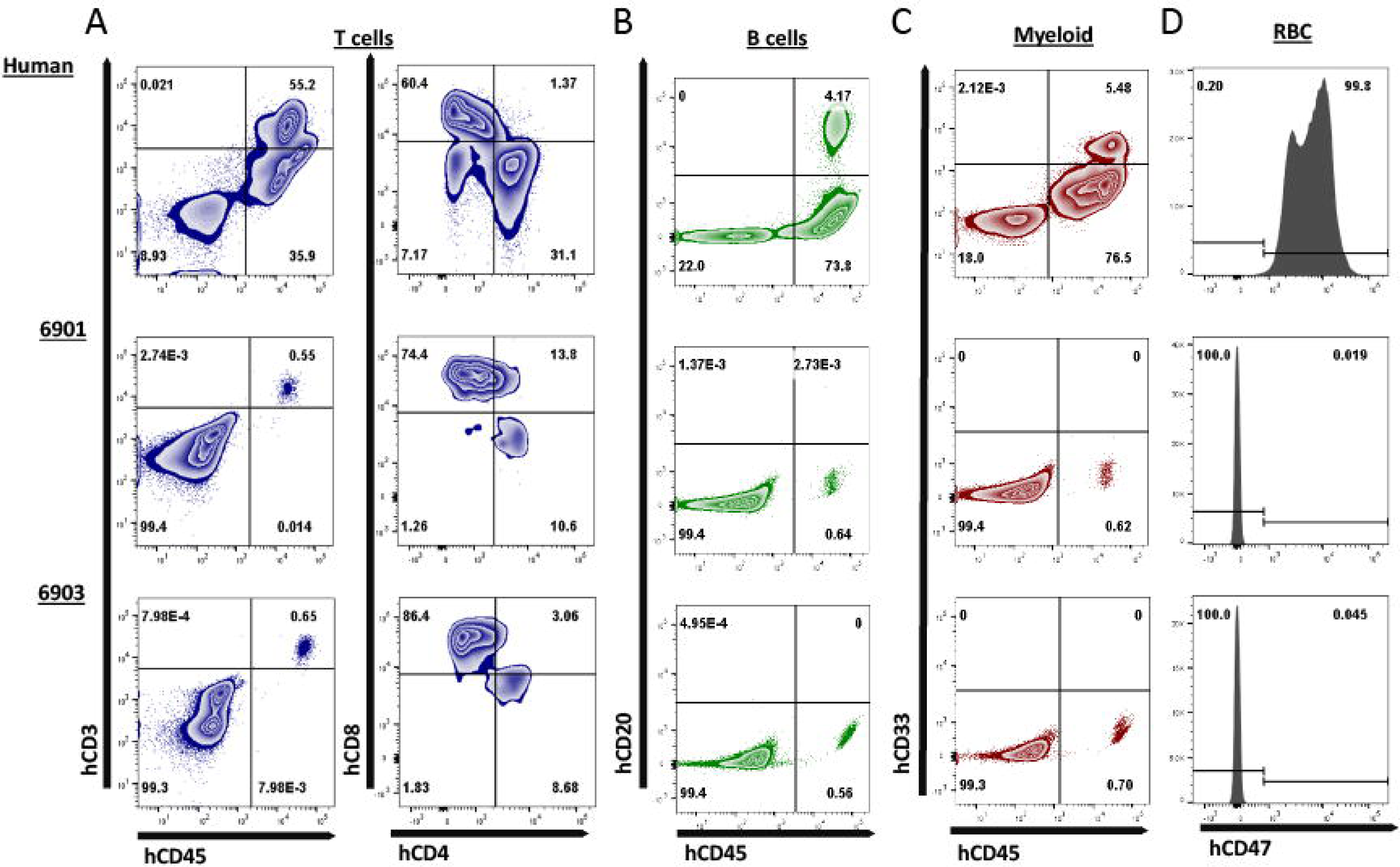
Human CD3ε+ cells are the primary cell type in peripheral blood of *Art*^-/-^ *IL2RG*^-/*Y*^ SCID pigs. Peripheral blood was collected from 6901 (day 0), 6902, and 6903 (Day 1) and stained for various human immune cell markers. Whole human blood was stained as a control to show gating strategy. All cells in circulation were gated from SCID pigs, while only lymphocytes and monocytes were gated from the human sample for the analysis of T, B, and myeloid cells. RBC were also stained separately for human CD47. **(A)** Presence of human T cells were assessed by staining for human CD45, CD3ε, CD4α, and CD8α. Cells that were CD3ε^+^ (left) were then assessed for CD4α and CD8α expression (right). Human T cells were present in both 6901 and 6903. Single positive CD4α^+^ and CD8α^+^ cells were also present. **(B)** Human B cells were assessed by staining for hCD45 and hCD20. B cells were not present in any pigs. **(C)** Human myeloid cells were assessed by staining for hCD45 and hCD33. Myeloid cells were not present in any pigs. **(D)** RBCs were stained for human CD47 expression to determine if human erythroid lineage differentiated in SCID pigs. No RBCs from SCID pigs expressed hCD47.

To create fetuses for human cell injections, one surrogate gilt was transplanted with 320 *Art*^-/-^ *IL2RG*^-/*Y*^ mutant embryos **(Supplemental Table 5)**. Pregnancy was confirmed at 28 days of gestation and laparotomy surgery for *in utero* injection was performed on gestational day 41. We injected half of the viable fetuses (n= 5 total; 3 injected) with 2 - 4 x 10^6^ cultured human CD34^+^ stem cells. The position in the uterine horn was recorded for all fetuses. The dam fully recovered from the laparotomy surgery and three piglets (6901, 6902, 6903) were delivered via Cesarean section at gestational day 119. The uterine positions at Cesarean section were noted and allowed identification of injected (6901, 6903) and un-injected (6902) piglets (see details in Methods). Cord blood was immediately collected from the three piglets to assess for the presence of human immune cells. We have tested and determined panels of anti-human antibodies that are not cross reactive to swine leukocytes **(Supplemental Fig. 5).** Whole cord blood was stained for pig and human CD45, as well as for human CD47. The two CD34^+^ cell injected pigs (6901 and 6903) were found to have human CD45^+^ (**Fig. 3B),** as well as human CD47^+^ cells in cord blood **(Fig. 3C).**

**Figure 5.**
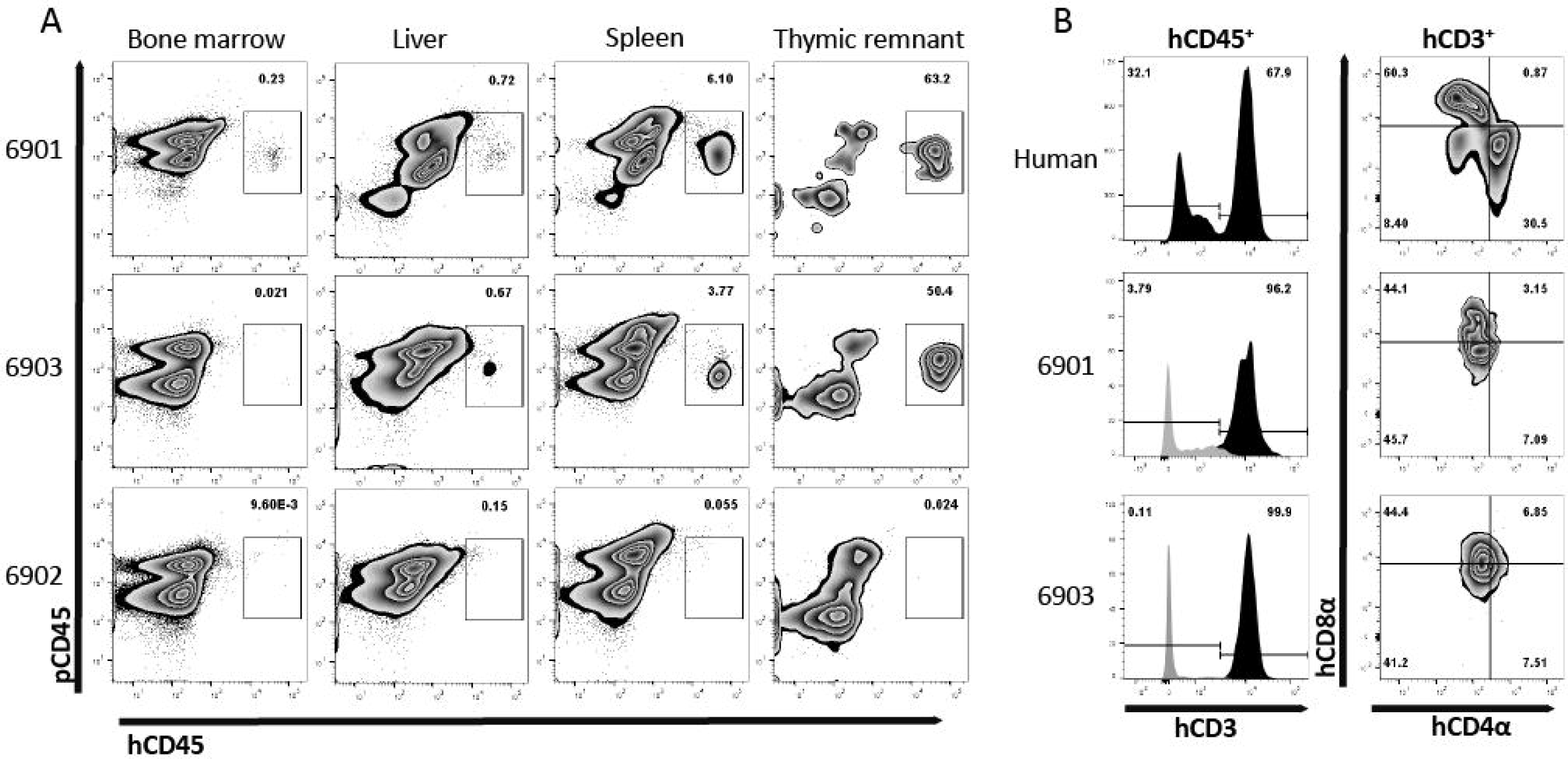
Human leukocyte engraftment in bone marrow, liver, spleen, and thymic tissue of *in utero* injected *Art* ^*-/-*^ *IL2RG* ^*-/Y*^ SCID pigs. **(A)** Lymphoid organs from 0-day old (6901) or 7-day old (6902 and 6903) SCID pigs were analyzed for the presence of human leukocytes by staining with human and pig CD45. Bone marrow liver, spleen, and thymic tissue from both 6901 and 6903 contained human CD45^+^ cells. All cells from isolated bone marrow and thymic tissue were stained, while mononuclear cells from spleen and liver were stained. **(B)** Human CD45^+^ cells in isolated cells from thymic tissue expressed human CD3ε. Black histogram is gated on hCD45^+^ cells, while grey is hCD45^-^ cells. Human CD4α and CD8α expression was assessed on human CD3ε^+^ cells within the thymus.

We next assessed if human cells were circulating in peripheral blood. One piglet (6901) was euthanized on day 0 due to a severe cleft palate. Peripheral blood was collected from 6901 at day 0 and at 1 day of age for 6902 and 6903. These blood samples were stained for human T cells (hCD3ε, hCD4α, and hCD8α) **(Fig. 4A),** B cells (hCD20) **(Fig. 4B),** and myeloid cells (hCD33) **(Fig. 4C).** Red blood cells were also stained for human CD47 **(Fig. 4D).** Nearly all human CD45^+^ cells circulating in 6901 and 6903 consisted of human CD3ε^+^ T cells that were positive for hCD4α and hCD8α (**Fig. 4A)**.

### Human CD45^+^ cell engraftment in Art^-/-^ IL2RG^-/Y^ bone marrow, liver, spleen, and thymic tissue

Since we observed human cells in peripheral blood, we next evaluated whether human immune cells were present within lymphoid organs. Interestingly, during necropsy, we observed grossly visible mesenteric lymph nodes in 6901 and 6903, but not in 6902. Additionally, all three pigs had some remnant, immature thymic tissue present over the heart **(Supplemental Fig. 6)**, which was collected for analysis. Of the tissues collected, cells were isolated from bone marrow, liver, spleen, and thymic tissue for flow cytometric analysis. Whole cell suspensions from bone marrow and thymus were stained, while mononuclear cells were stained from liver and spleen. Human CD45^+^ cells were found in all four tissues assessed in 6901 and 6903, which both had human CD45^+^ cells in circulation **(Fig. 5A).** At least half of the isolated thymic tissue cells from these two animals were human cells. Animal 6902 did not have any human cells in any lymphoid organs.

**Figure 6.**
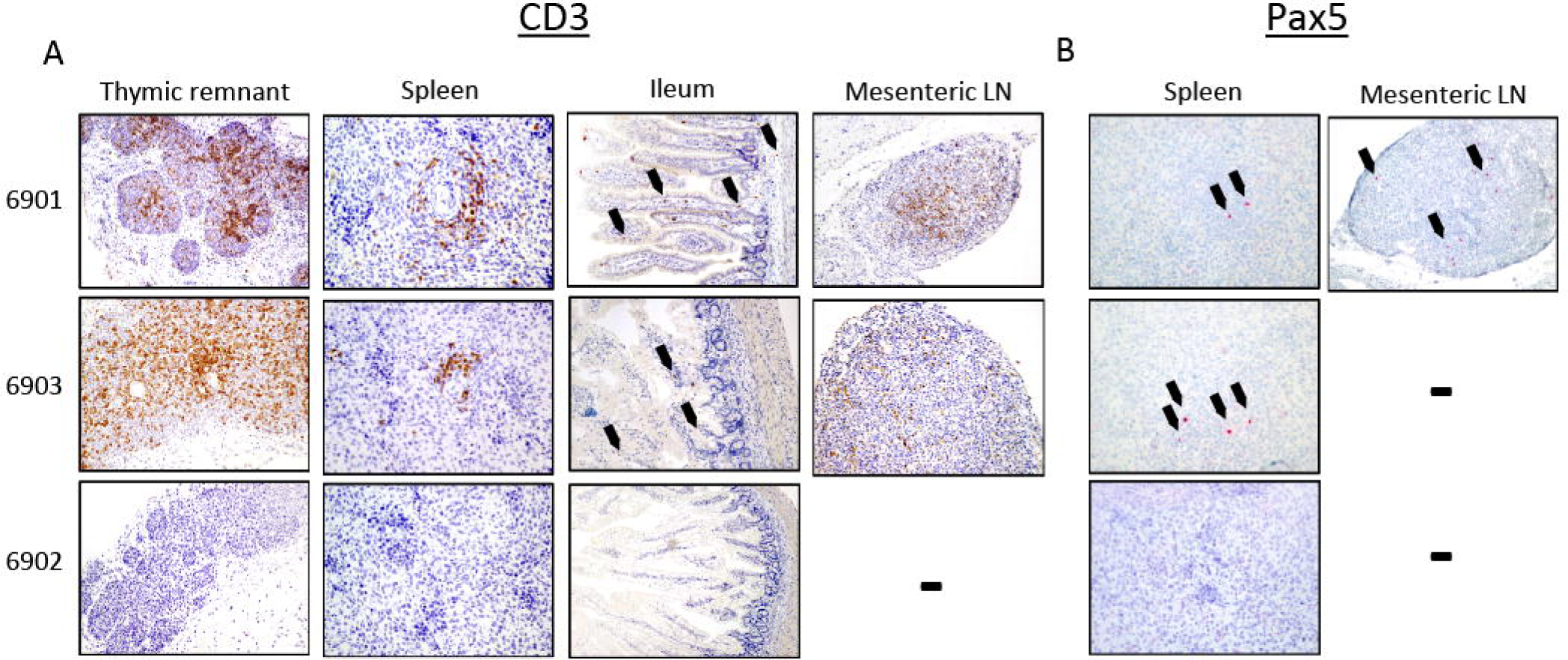
Human CD3ε^+^ and Pax5^+^ cells in lymphoid organs of *in utero* injected *Art* ^*-/-*^ *IL2RG* ^*-/Y*^ SCID pigs. Lymphoid tissues were collected and stained for **(A)** CD3ε and **(B)** Pax5. **(A)** Thymic tissues from 6901 and 6903 were robustly populated with CD3ε^+^ cells, and CD3ε^+^ cells were also present in periarteriolar sheaths within the spleen. The ileum tissue from both animals had punctate CD3ε^+^ cells (black arrows); CD3ε^+^ cells were also present within mesenteric lymph nodes (LN). Lymphoid tissues from 6902 did not contain CD3ε^+^ cells. **(B)** Spleen from both 6901 and 6903 had punctate Pax5^+^ cells. Only the mesenteric LN in 6901 had Pax5^+^ cells. No Pax5^+^ cells were found in tissues from 6902.

Since a majority of the human cells we observed in these pigs were hCD3ε^+^, we assessed the expression of hCD4α and hCD8α within the thymic tissue cells. Early in development, T cells express both CD4α and CD8α. In animal 6901, we observed that the thymic hCD45^+^ hCD3ε^+^ cells were either hCD4α^+^ hCD8α^-^ or hCD4α^-^ hCD8α^-^, while in 6903 they appeared to be hCD4α^-^ hCD8α^-^ **(Fig. 5B)**.

In addition to flow cytometric analysis of cells within lymphoid tissues, we also analyzed tissues by IHC. Lymphoid tissues were collected and assessed for the presence of human T and B cells in the *in utero* injected piglets. Thymic tissue, spleen, ileum, and mesenteric lymph nodes from both 6901 and 6903 had CD3ε^+^ cells **(Fig. 6A)**. Spleens from both 6901 and 6903 also had punctate Pax5^+^ cells present, which is a marker for B cell development. A mesenteric lymph node from 6901 also had Pax5^+^ cells **(Fig. 6B)**. Tissues from 6902 did not stain positively for either CD3ε or Pax5. These histology results are consistent with the flow cytometric analyses demonstrating human leukocyte engraftment within *Art*^-/-^ *IL2RG*^-/*Y*^ SCID pigs injected with human CD34^+^ cells.

## Discussion

Herein we have described foundational steps towards the development of an immunologically humanized large animal SCID model. To create the model, we introduced a mutation into the *IL2RG* gene using the CRISPR/Cas9 system in a naturally occurring *Art*^-/-^ SCID background to generate *Art*^-/-^ *IL2RG*^-/*Y*^ pigs that lacked T, B, and NK cells. We performed a pig to pig BMT procedure on one male *Art*^-/-^ *IL2RG*^-/*Y*^ pig, which led to successful reconstitution of graft T and NK cells, but very few B cells. Next, we utilized *in utero* injection procedures to introduce human hematopoietic stem cells into the intraperitoneal space of SCID pig fetuses. We observed that human CD3ε^+^ cells were present in bone marrow, spleen, liver, and thymic tissue in injected pigs after birth. Human Pax5^+^ cells were also present within the spleen and mesenteric lymph nodes of injected pigs. Intrahepatic injection of human CD34+ cells in newborn *NOD Rag*^-/-^ *IL2RG*^-/-^ mice has resulted in similar patterns of reconstitution (T and B cell development) as our *Art*^-/-^ *IL2RG*^-/*Y*^ pig ^46^.

### Improving human cell engraftment in SCID pigs

One surprising finding was that we did not detect human myeloid cells, as has previously been reported in past *in utero* injections of human CD34^+^ cells in immunocompetent pig fetuses^47^. In this initial humanization model, we had a pre-determined end point of 7 days to assess human cells in blood and tissues, and therefore we only probed for myeloid cells during this period. In previous studies, myeloid lineage development had been assessed at 40 days post injection (80 days of gestation)^41^. It may be that human myeloid cells are transient during gestation in this fetal injection model. Further investigation is needed to improve human myeloid reconstitution in neonatal *Art*^-/-^ *IL2RG*^-/*Y*^ SCID pigs.

In the process of *in vitro* CD34^+^ cell culture with SCF, TPO, and FLT-3L, some cells may lose their stemness, which likely contributed lack of a variety of cells that differentiated (i.e. only T and B cells) within our SCID pig model. In mouse models genetic modifications have been required to attain human myeloid and NK cell engraftment, including human CSF-1, IL-15, GM-CSF, Flt-3L, IL-3, TPO, as human cells do not recognize mouse cytokines^21,23,47–50^. An area to be investigated is how human cells respond to swine cytokines and if the swine bone marrow niche is supportive of human myeloid cells. Humanization of certain cytokine genes may be required in future humanization attempts in our pig model. *In vitro* culturing assays with human hematopoietic stem cells and porcine cytokines can be a first-line screen for assessing porcine cytokine cross-reactivity. Additionally, in future studies, we can assess cytokine secretion by developed human cells within the pigs.

In our model, we observed that a majority of human cells that developed were CD3ε^+^. We assessed the thymic tissue to better understand the development of human T cells. We interestingly did not observe CD4α^+^CD8β^+^ double positive cells within the thymus, which is an expected normal stage of T cell development. In future studies, it will be imperative that we perform deeper phenotyping of human cells that have differentiated within the pig. Previous reports by Kalscheuer *et al*.^51^ and Ogle *et al*.^42^ show that the swine thymus can support engraftment and differentiation of human T cells. The lack thymic development in a SCID pig fetus may negatively impact the ability of human cells to develop, which may warrant transplantation of human thymic tissue after birth.

Another potential method to increase engraftment is to condition the fetuses prior to human cell injection. Plerixafor, a drug that mobilizes stem cells out of bone marrow^52^, has previously been utilized for *in utero* injections of human cells into sheep fetuses to improve engraftment of human cells^53^. Plerixafor is an agonist for CXCL12 on stromal cells, which binds to CXCR4 on hematopoietic stem cells (HSC)^52^. Administration of plerixafor mobilizes sheep HSC out of bone marrow, providing more available niches for human HSC to engraft. Goodrich et al. (2014) described that administration of plerixafor along with injection of CD34^+^ CXCR4^+^ human stem cells and mesenchymal stem cells improved chimerism (in peripheral blood) 5 weeks after transplantation from 2.80 to 8.77%. Now that T^-^ B^-^ NK^-^ SCID pigs and biocontainment facilities are available for extended postnatal follow-up, a similar regimen could be administered to SCID pig fetuses prior to *in utero* injection with human stem cells.

### Outlook on B cell reconstitution in Art^-/-^ ILR2G^-/Y^ SCID pigs

One issue in both pig to pig BMT, as well as *in utero* injection of human HSCs, was the failure of pig or human B cells to robustly develop. Historically, some human SCID patients that underwent BMT have also failed to develop graft derived B cells^33^. In some cases, significant B cell reconstitution in human BMT can require up to two years ^54,55^. One leading hypothesis regarding B cell development issues is due to differences in the B cell niche in the bone marrow. Single mutant *IL2RG* knock out pigs are capable of developing B cells, which can be detected in circulation^9,10^, however they are non-functional due to the absence of T helper cells. Mutations in *Artemis* lead to a B cell block of differentiation at the pre-B cell phase^56^. Together, an *Art*^-/-^ *Il2RG*^-/*Y*^ pig likely still has premature B cells present in the bone marrow, which would prevent further engraftment and differentiation of graft stem cells in this niche. Conditioning prior to stem cell transplantation has helped improve B cell reconstitution in some cases^57^, although such conditioning procedures would be difficult prior to *in utero* cell transplantation. To our knowledge this is the first time a bone marrow transplantation has been performed on a double mutant SCID pig. Thus, further assessment of the bone marrow niches of Art Il2RG pigs may be required to better understand conditioning regimens that may be needed for engraftment.

### Concluding remarks

As the field of biomedical SCID pig research expands, new techniques will arise to optimize human cell engraftment within SCID pig models. We can draw from previous large animal *in utero* injection protocols^41–44,53,58^, as well as humanization techniques performed in immunocompromised mice ^59,60^. In our SCID pig model, we show that human T and B cells can develop. As we improve reconstitution of human cell subsets, the humanized SCID pig will be a critical alternative large animal model for researchers preclinical or co-clinical trials.

## Materials and Methods

### Study design

Our study was designed to develop a T^-^ B^-^ NK^-^ SCID pig model by generating *Art*^-/-^ *IL2RG*^-/*Y*^ pigs by CRISPR Cas9 site directed mutagenesis of our existing *Art*^-/-^ pig line, which we discovered in 2012^1,2,61^. We aimed to generate these pigs as a large animal biomedical model for human cell and tissue xenotransplantation. Once we successfully created the *Art*^-/-^ *IL2RG*^-/*Y*^ fibroblast cell line, we performed a total of 8 embryo transfer surgeries to generate piglets. Of these transfers, 5 females became pregnant, and a total of three litters were born; one of which we performed *in utero* injections of human CD34^+^ cells. Once piglets were born, we confirmed their T^-^ B^-^ NK^-^ phenotype. We performed a pig to pig bone marrow transplant on one *Art*^-/-^ *IL2RG*^-/*Y*^ boar, which would allow us to eventually collect semen for future breeding and use of this genetic line. We performed *in utero* injections of human cord blood selected CD34^+^ cells on *Art*^-/-^ *IL2RG*^-/*Y*^ fetuses from one pregnant female. Three piglets were born from this litter, with two piglets showing evidence of human immune cell engraftment. The low number of animals in this study are a result of small litter sizes of cloned piglets, as well as low pregnancy rates of embryo transfer procedures.

### Ethics statement

All animal protocols were approved by Iowa State University’s Institutional Animal Care and Use Committee. All animals were utilized in accordance with the Animal Welfare Act and the Guide for the Care and Use of Laboratory Animals. All human sample collection protocols were approved by Iowa State University’s Institutional Review Board.

### Establishment of porcine fetal fibroblast cell lines with Art^-/-^ genetic background

Our population of SCID pigs has two natural mutations in two separate *Artemis* alleles, termed *Art12* and *Art16*. The *Art12* allele contains a nonsense mutation in exon 10, while the *Art16* allele contains a splice site mutation in intron 8^3^. Frozen semen from a bone marrow transplanted (BMT) rescued *Art12*/*16* ^32^ boar was utilized to artificially inseminate two *Art*^+/-^ carrier sows. The sows were sacrificed at day 35 of gestation and fetuses were collected in a sterile manner to obtain fetal fibroblasts (pFF), as described previously^62^. Briefly, minced tissue from each fetus was digested in 20 mL of digestion media (Dulbecco-modified Eagle medium [DMEM] containing L-glutamine and 1 g/L D-glucose [Cellgro] supplemented with 200 units/mL collagenase and 25 Kunitz units/mL DNaseI) for 5 h at 38.5°C. After digestion, pFF cells were washed in sterile PBS and cultured in DMEM supplemented with 15% fetal bovine serum (FBS) and 40 μg/mL gentamicin. Upon reaching 100% confluence, the pFF cells were trypsinized, frozen in FBS with 10% dimethyl sulfoxide (DMSO) and stored long-term in liquid nitrogen. Simultaneously, cellular DNA was sent for SLA typing, as described in Powell et al^32^. *SRY* and *Artemis* primers **(Supplemental Table 6)** were utilized to identify pFF sex and *Artemis* genotype^3^.

### CRISPR/Cas plasmid and sgRNA product

Guide RNAs targeting exon 5 of *IL2RG* were designed utilizing software available from Zhang Lab (https://zlab.bio/guide-design-resources). The sequence of the designed sgRNA was: 5’-GGCCACTATCTATTCTCTGA**AGG**-3’; the bold font identifies the PAM site. The sgRNA oligos were annealed and ligated into the human codon-optimized SpCas9 expression plasmid (pX330; Addgene plasmid # 42230), as described previously^63^. We only transfected male *Art12*/*12* cell lines (herein referred to as *Art*^-/-^).

### Identification of off-target sequences

To identify putative off-target sequences for the CRISPR/Cas9 mutagenesis used in *Art*^-/-^ *IL2RG*^-/*Y*^ piglets, bioinformatics tools (http://www.rgenome.net/cas-offinder/) were used. Ten potential off-target sites were identified and primers for the off-target positions were designed. Genomic DNAs obtained from ear notches of *Art*^-/-^ *IL2RG*^-/*Y*^ pigs were used as templates in PCR amplification of potential off-target regions. Primers and gene information for this purpose are in **Supplemental Table 7**. DNA sequencing results revealed no mutations had occurred in any of the potential off-target positions.

### Establishment of transfected clonal colonies and identification of IL2RG mutagenesis

Male Art^-/-^ pFFs were used for cell transfection, as described in Whitworth et al^64^. Briefly, pFFs were cultured in 75cm^2^ flasks to reach 90% confluency, trypsinized, resuspended at a concentration of 1.0 x 10^6^ cells/mL in Electroporation Buffer medium (25% Opti-MEM and 75% cytosalts [120 mM KCl, 0.15 mM CaCl2, 10 mM K2HPO4; pH 7.6, 5 mM MgCl2]), and prepared for transfection. A mixture of 1 μg sgRNA ligated vector and 200 μL of cell suspension in electroporation buffer was then transferred into 2 mm gap cuvettes (Fisher Sci, 9104-6050) and exposed to three-1 msec square-wave pulses at 250 V, using the BTX Electro Cell Manipulator (Harvard Apparatus). Transfected cells were then diluted, 80-200 cells were plated in 100 mm culture dishes to obtain distinct clonal cell colonies and maintained at 38.5°C in 5% CO_2_. After 10-12 days in culture, cell colonies were delineated using cloning cylinders and picked for clonal colony propagation and DNA sequencing. Primers **(Supplemental Table 6)** flanking the *IL2RG* exon 5 target region were utilized to test clonal colonies and PCR products thus obtained were purified by ExoSAP-IT PCR product Cleanup kit (Affymetrix Inc, Thermo Fisher Scientific). Mutant PCR products were cloned into PCR2.1 vectors (Life Technologies) and transformed into *E.coli* DH5-α maximum competent cells (Life Technologies). Ten colonies were chosen and DNA from these samples were sent to the Iowa State University DNA Facility for sequencing. Sequences were aligned by Bio-Edit software (Ibis Biosciences, Carlsbad, CA, USA) for comparison with wild-type alleles, to identify cell lines with the appropriate mutations in *IL2RG* on the *Art*^-/-^ background.

### Double mutant embryo production and surgical embryo transfer

Purchased pig oocytes (DeSoto Biosciences, Inc.) or those derived from aspirating ovaries collected from a local abattoir were utilized for *in vitro* maturation (IVM), as previously described ^65,66^. Briefly, oocytes were matured *in vitro* with maturation medium (TCM-199 with 2.9 mM HEPES, 5 μg/mL insulin, 10 ng/mL epidermal growth factor, 0.5 μg/mL porcine follicle stimulating hormone, 0.91 mM pyruvate, 0.5 mM cysteine, 10% porcine follicular fluid, and 25 ng/mL gentamicin), and transferred into fresh medium after 22 h. Following IVM, cumulus-oocyte-complexes (COC) were vortexed for 3 minutes in 0.1 % hyaluronidase in TCM199 with HEPES to obtain denuded oocytes. Metaphase II (MII) oocytes, identified by the presence of an extruded polar body, were placed in manipulation medium (TCM199 with HEPES supplemented with 7 μg/mL cytochalasin B) and used thereafter for somatic cell nuclear transfer (SCNT). The extruded polar body, along with a portion of the adjacent cytoplasm, presumably containing the M II plate, were removed, and a donor nucleus of the appropriate *Art*^-/-^ *IL2RG*^-/*Y*^ genotype was placed in the perivitelline space by using a thin glass capillary. The reconstructed embryos were then placed in a fusion medium (0.3 M mannitol, 0.1 mM CaCl_2_, 0.1 mM MgCl_2_, and 0.5 mM HEPES) and exposed to two DC pulses (1-sec interval) at 1.2 kV/cm for 30 μsec using a BTX Electro Cell Manipulator (Harvard Apparatus). After fusion, these embryos were activated in embryo activation medium (10 μg/mL cytochalasin B) for 4 h. After chemical activation, cloned zygotes were treated with 500 mM Scriptaid for 12-14 h and then cultured in porcine zygote medium 3 (PZM-3) until embryo transfer. Embryos produced over two days were then surgically transferred into the ampullary-isthmic junction of the oviduct of the surrogate on day 1-2 post estrus. Pregnancies were confirmed by ultrasound approximately 30 days following embryo transfer.

### Cesarean section and rearing of Art^-/-^IL2RG^-/Y^ SCID pigs

At gestational day 119, pregnant gilts underwent Cesarean sections. We chose gestational day 119 instead of 114 (normal gestation) because piglets derived from somatic cell nuclear transfer typically requiring longer gestational period. Initial anesthesia was induced with either a lumbar epidural of propofol (0.83-1.66 mg/kg) or intravenous injection of Ketamine (1-2 mg/kg) and Xylazine (1-2 mg/kg), and anesthesia was maintained on oxygen and isoflurane. An abdominal incision was made to expose and remove the uterus. After removal, the uterus was immediately rinsed in chlorohexidine and then surgically opened to remove the piglets. All piglets had their cords clamped before being immediately placed into sterile polystyrene boxes and delivered into biocontainment facilities^14^. All piglets were fed approximately 250 mL of pasteurized porcine colostrum within the first 24 hours of life. Piglets derived from the gilt that underwent laparotomy procedures for human stem cell injection (see below) did not receive colostrum.

### Genotyping of neonatal Art^-/-^ IL2RG^-/Y^ pigs

After birth, DNA was isolated from ear notch tissues and subjected to genotyping for *Art* and *IL2RG* status using primers and protocols described in **Supplemental Table 6.**

### Flow cytometry staining for Art^-/-^ IL2RG^-/Y^ pig characterization and bone marrow engraftment monitoring

Whole blood or cord blood from newborn *Art*^-/-^ *IL2RG*^-/*Y*^ piglets was collected into an EDTA blood collection tube. Whole blood was stained for porcine CD3ε, CD8α, and CD172α to assess the presence of porcine T and NK cells. Cells were additionally stained for porcine CD21 to assess the presence of B cells **(Supplemental Fig. 1)**. Blood from the *Art*^-/-^ *IL2RG*^-/*Y*^ BMT was also stained for CD79 to assess B cell reconstitution **(Supplemental Fig. 3)**. Additional information about the antibodies used can be found in **Table 1**.

Blood was collected from the bone marrow transplanted boar approximately once a month after the BMT and subjected to either a complete blood count (CBC) at Iowa State University’s Veterinary Diagnostic lab or by flow cytometry analysis using the above listed antibodies. All samples were run on a custom BD LSR II and data were analyzed using Flowjo (Tree Star).

### Pig bone marrow isolation and bone marrow transplantation

A complete SLA-matched female sow of approximately 4 years of age was euthanized and used as a bone marrow donor for one *Art*^-/-^ *IL2RG*^-/*Y*^ piglet. Briefly, sternum and ribs collected from the animal were dipped in 70% ethanol after collection. A sterilized Dremel tool was used to make holes halfway through the bone to expose bone marrow. Sterilized Spratt Brun bone curettes were used to scrape marrow from the bone. HBSS (without phenol red) was used to flush the bone marrow to collect cells; any other loosened marrow was also placed in HBSS. After marrow isolation, the suspension was washed in HBSS. The cells were resuspended in ACK lysing solution (Lonza) for ten minutes at room temperature. The suspension was washed in HBSS and filtered through a 70 μm cell strainer. A total of 2.27 × 10^8^ million unfractionated cells were isolated and resuspended in approximately 3 mL of HBSS for infusion.

To infuse bone marrow-derived cells into the *Art*^-/-^ *IL2RG*^-/*Y*^ recipient, the five-day old SCID piglet was anesthetized with and maintained on isoflurane gas during the procedure. A catheter was placed in an ear vein and the cell suspension slowly infused. After infusion, personnel monitored the piglet until fully recovered.

### Immunohistochemistry of human and porcine immune markers

Lymphoid organs were collected into 3.7% formaldehyde in 1X PBS for 24 hours. Tissues were then moved to 70% ethanol until processing. IHC staining for T and B lymphocyte markers (for *Art*^-/-^ *IL2RG*^-/*Y*^ immune characterization) was performed in paraffin-embedded tissue thin sections at the Kansas State Veterinary Diagnostic Laboratory (KSVDL). Briefly, deparaffinized slide-mounted thin sections were pre-treated for 5 minutes with a peroxide block, followed by incubation with primary antibody. A mouse monoclonal anti-CD3ε (clone LN10, Leica Biosystems) was used to stain for pig T cells, while a mouse monoclonal anti-CD79α (clone HM57, Abcam) was used to stain for B cells. Primary antibodies were incubated with PowerVision Poly-HRP anti-mouse IgG at room temperature for 25 minutes with DAB chromagen, and then counterstained with hematoxylin.

Lymphoid tissues from SCID pigs engrafted with human cells were treated similarly as above. Staining was performed at Michigan State University’s Department of Pathobiology and Diagnostic Investigation. Tissues were stained for anti-CD3ε (Dako #A0452) and anti-Pax5 (Ventana clone 24) to assess for the presence of human T and B cells, respectively. Briefly, CD3ε staining was performed by antigen retrieval with standard ER1 retrieval for 20 minutes, and tissues were analyzed on a BondMax for the detection of DAB chromagen. For Pax5 staining, antigen retrieval was performed with standard CC1 retrieval for 64 minutes, and tissues were analyzed on a Discovery Ultra AP for ultrared detection.

### Human hematopoietic stem cell isolation from cord blood

Human cord blood was collected at the Mary Greeley Medical Center in Ames, Iowa, into 50 mL conical tubes containing 8 mL of anticoagulant citrate dextrose solution (38 mM citric acid, 85.25 mM sodium citrate, 136 mM dextrose). Mononuclear cells (MNCs) were isolated from cord blood by diluting blood 1:2 in HBSS and then layering over Ficoll-Paque. Buffy coats were collected and washed in HBSS. Prior to stem cell isolation, MNCs were resuspended in an isolation HBSS (iHBSS) consisting of 0.5% FBS and 2 mM EDTA in HBSS.

To isolate human CD34^+^ cells, we used a CD34 MicroBead kit from Miltenyi Biotech. Briefly, MNCs were incubated in iHBSS with CD34 microbeads and FcR blocking reagent for 30 minutes at 4° C on a rocker. After the incubation period, cells were washed in iHBSS and then passed through a LS column in a Miltenyi magnet to capture human CD34^+^ cells. The column was then removed from the magnet and cells were flushed, washed once more in HBSS, and then frozen in 10% DMSO and 90% FBS at −80° C until use.

### Human HSC thawing, culturing, and preparation for fetal injection

Human CD34^+^ cells were thawed by diluting into complete RPMI media (10% FBS, 2 mM glutamine, 50 μg/mL gentamicin, and 10 mM HEPES). Cells were cultured in Miltenyi StemMACS media containing Thrombopoietin (TPO), Stem Cell Factor (SCF), and Flt3-Ligand (FLT-3L), at starting concentrations of 38-42 × 10^3^ cells/mL. Cells were left in culture for 7 days and expanded 184.5-fold. Cells were prepared for injection by washing three times in phosphate buffered saline and resuspended at a concentration of 26.6 × 10^6^ cells/mL. A total of 150 μL of cell suspension with either 2 or 4 × 10^6^ cells were administered to the fetuses in 0.9% saline.

### Laparotomy procedure for fetal injection of human stem cells

Laparotomy procedures were performed as previously described in Boettcher *et al*. (2019)^45^. Briefly, the gilt was started on 15 mg of Matrix orally one day before surgery and maintained on Matrix until gestational day 118. Immediately prior to sedation, the gilt was given 0.01 mg/kg Glycopyrrolate by intramuscular injection and then anesthetized with 2 mg/kg Xylazine and 5 mg/kg Telazol by intramuscular injection. The gilt was then placed in dorsal recumbency, intubated, and started on isoflurane (3-5%) and oxygen (2.5 L/min). Lactated Ringer’s solution was given in an ear catheter at a constant rate infusion within 10 minutes of the first incision. The abdominal area was scrubbed with chlorohexidine and the surgical field was covered with sterile drapes and Ioban drapes.

A ventral midline incision was made from the caudal most nipple extending to the caudal aspect of the umbilicus through the *linea alba* into the peritoneal cavity. The left uterine horn was exposed and visualized with a ZONARE ultrasound with an L14-5sp intraoperative linear array transducer (10 MHz). Two live and one nonviable fetus were visualized in the left horn; one was injected with 4 million cultured human stem cells in a volume of 0.15 mL within the intraperitoneal space. The left horn was placed back into the abdominal cavity and the right horn was exposed and visualized, and three fetuses were observed. One fetus was injected with 2 million and another fetus with 4 million human stem cells within the intraperitoneal space. In total, three out of the five viable fetuses were injected. The abdominal cavity was lavaged with 500 mL of Lactated Ringer’s solution and sutured closed. The gilt then received 0.18 mg/kg Buprenophine-Sustained Release subcutaneously, 5 mg/kg Ceftiofur Crystalline Free Acid (Excede), and 0.3 mg/kg of meloxicam and monitored by personnel until recovered from anesthesia. The gilt recovered from the laparotomy surgery with no issues and underwent a Cesarean section at day 119 of gestation, as described above.

At the time of Cesarean section, we observed two live piglets within the left horn and one live piglet in the right horn of the uterus. During the laparotomy, we observed two viable fetuses in the left horn, of which one was injected, and their relative position was recorded. After Cesarean delivery, by position in the uterus we were able to identify the two developed piglets in the left horn as injected (6901) and non-injected (6902). The right horn originally had three viable fetuses but only one piglet survived to term. After flow cytometric analysis, we confirmed that the one piglet on the right horn (6903) had been injected based on presence of human cells.

### Isolation of MNC from tissue for flow cytometric analysis

To collect lymphoid MNCs, tissue was collected and placed into Hanks Balanced Salt Solution (HBSS) with 10μg/mL gentamicin. Tissues were minced in a digestion solution of HBSS with 300 mg/mL of collagenase, 3% FBS, and 2mM HEPES (referred to as d-HBSS). The tissue incubated for 1 hr at 37°C with vortexing every 15 minutes, and then strained over a 70 μm cell strainer. Cells were washed once and counted with a BD counting kit, and then were stained as described below.

### Flow cytometry analysis for human cells in Art^-/-^ IL2RG^-/Y^ pig blood

Peripheral or cord blood was collected from piglets into EDTA blood containers. Whole blood or MNC from tissues were stained with antibodies against human and pig cell subset markers **(Table 1)**. Each staining step was incubated for 15 minutes at 4°C and washed with 1X PBS with 0.1% sodium azide. Red blood cells were lysed with ammonium chloride lysing solution. Cells were fixed in 2% formaldehyde in 1X PBS and data were acquired on a FACs Canto II flow cytometer and analyzed using FlowJo (Treestar).

## Acknowledgements

We thank Iowa State University’s animal care staff for care of SCID animals throughout the study.

## Author Contributions

ANB designed humanization experiments, isolated human stem cells, performed fetal injections, human immune cell flow cytometry, and wrote the draft of the manuscript. YL isolated fetal fibroblasts, performed CRISPR/Cas9 genome modification, performed somatic cell nuclear transfer and embryo transfers to generate double mutant piglets, and wrote the draft of the manuscript. AA performed fetal injection laparotomy procedures on pregnant gilt. MK performed immunohistochemistry for human immune cells in lymphoid tissues. KB and CL performed flow cytometry for *Art*^-/-^ *IL2RG*^-/*Y*^ immunophenotyping and bone marrow transplantation monitoring. AGCO and JW performed immunohistochemistry for porcine immune cells in lymphoid tissues. EP performed IV injections of human stem cells in *Art*^-/-^ pigs. JS collected cord blood for stem cell isolations that were used in fetal injections of *Art*^-/-^ *IL2RG*^-/*Y*^ fetuses. ES performed pig bone marrow isolation for bone marrow transplantation. CSH performed MHC PCR analysis for bone marrow donors. JWR performed bone marrow transplantation. SC genotyped fetal fibroblast cell lines for *Art* status. SC, ZK, and MA were involved in experimental planning for all facets of this project. JC was involved in experimental flow cytometry planning. SS performed numerous ultrasound pregnancy checks on surgically transferred gilts. GD and JJ assisted with fetal injection procedures. FG and EW provided human stem cells for IV injection procedures in *Art*^-/-^ pigs. JCMD provided guidance in pig breeding to generate the litters for pFF collection and assisted in project planning discussions. JWR and CKT were involved in all aspects of procedures and experiments performed in this study and edited the manuscript.

