## Supplemental Information for "Novel engraftment and T cell differentiation of human hematopoietic cells in *Art*^-/-^ *IL2RG*^-/^ SCID pigs"

**List of Supplementary Figures**

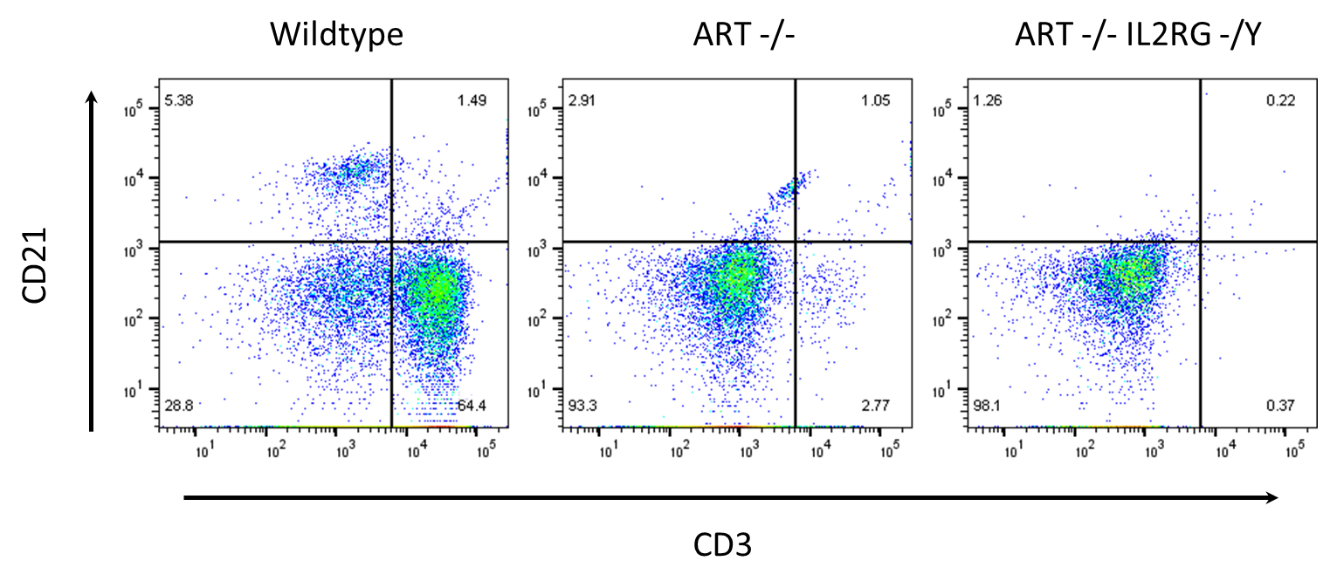

**Supplemental Figure 1. B cells are absent in *Art^-/-^ IL2RG^-/Y^* pigs.** Whole blood was stained for CD3ε and CD21 from a wildtype, *Art^-/-^* and *Art^-/-^ IL2RG^-/Y^* pigs. *Art^-/-^* pigs lack B cells, although they have a “leaky” T cell phenotype. *Art^-/-^ IL2RG^-/Y^* pigs do not have T or B cells in circulation.

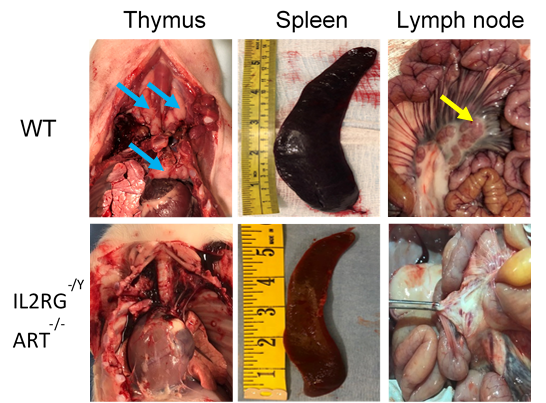

**Supplemental Figure 2. Lymphoid organs from wildtype and *Art^-/-^ IL2RG^-/Y^* pigs.** *Art^-/-^ IL2RG^-/Y^* pigs lack a thymus, have an atrophied spleen, and also lack mesenteric lymph nodes. Blue arrows indicate thymus. Yellow arrow indicates mesenteric lymph nodes.

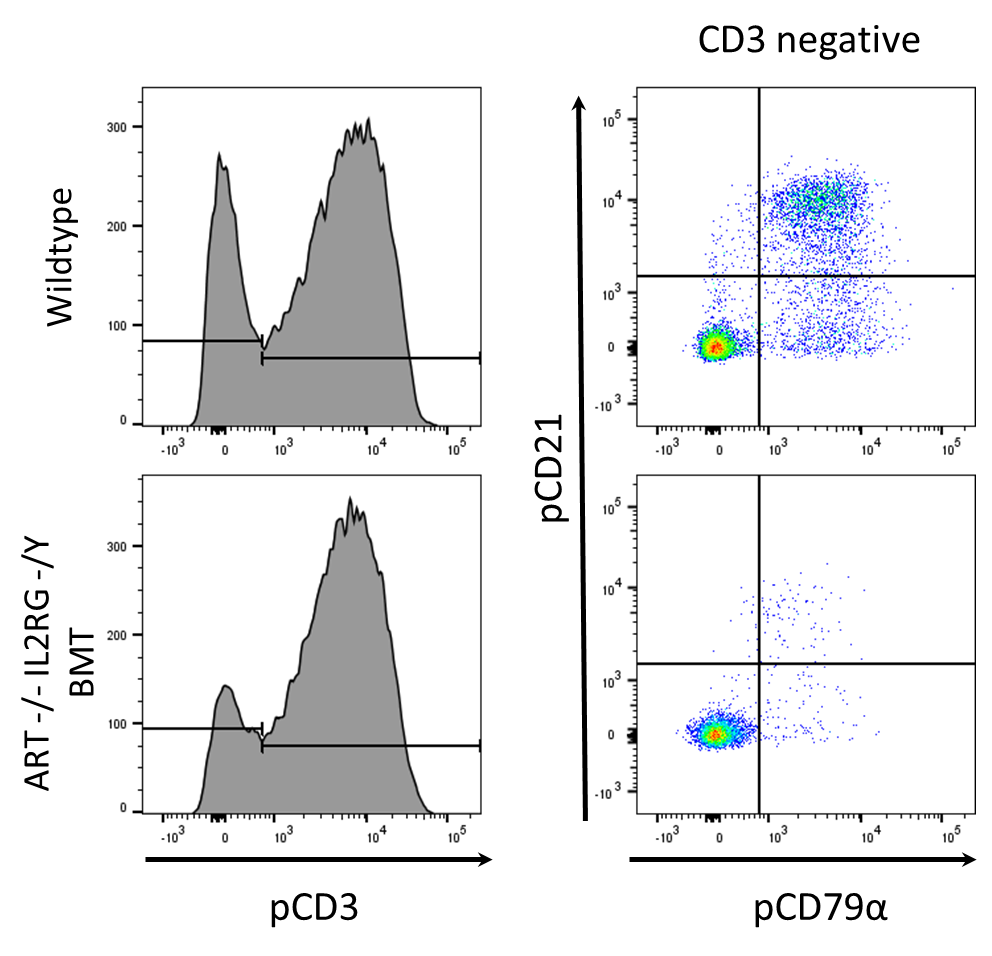

**Supplemental Figure 3. Decreased engraftment and differentiation of B cells in an *Art^-/-^ IL2RG^-/Y^* SCID pig 5 months post bone marrow transplant.** Whole blood was collected from a wildtype and BMT *Art^-/-^ IL2RG^-/Y^* SCID pig and stained for CD3ε, CD79α, and CD21. Lymphocytes were gated and first assessed for expression of CD3ε. The CD3ε negative population was then assessed for expression of CD21 and CD79α. The BMT SCID pig has normal proportions of T cells, while lacking a B cell population.

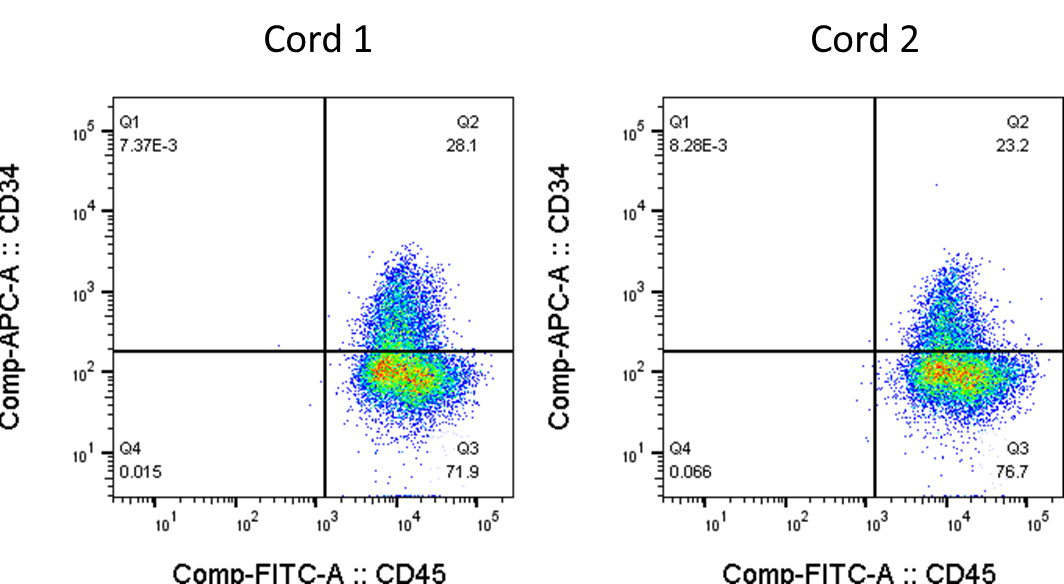

**Supplemental Figure 4. Human CD45 and CD34 expression on *in utero* injected stem cells.** Human CD34^+^ stem cells were isolated from cord blood and cultured for 7 days prior to *in utero* injections with FLT-3L, stem cell factor, and thrombopoietin. Piglet 6901 was injected with cultured stem cells from Cord 1, while piglet 6903 was injected with stem cells from either Cord 1 or Cord 2.

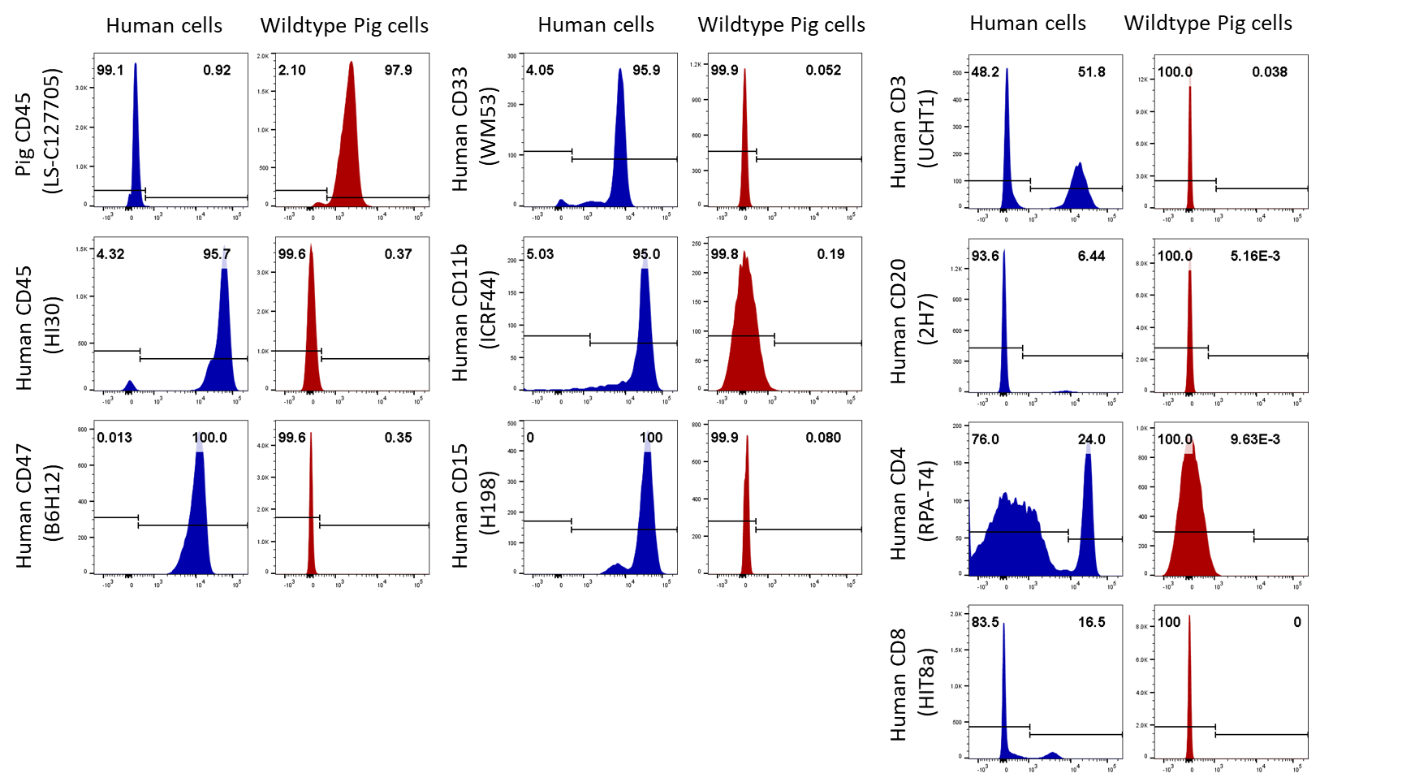

**Supplemental Figure 5. Species specific antibodies utilized to detect human cells in *in utero* injected SCID pigs.** Whole blood was collected from a human and a wild type pig and stained for a variety of markers found in lymphoid, myeloid, and granulocytes. Markers and clones are labeled above; specific antibody information can be found in **Table 1**. All cells were gated for the analysis of pCD45, hCD45, and hCD47. Only monocytes were gated for the analysis of hCD33 and hCD11b. Only granulocytes were gated for the analysis of hCD15. Only lymphocytes were gated for the analysis of hCD3ε, hCD20, hCD4α, hCD8α.

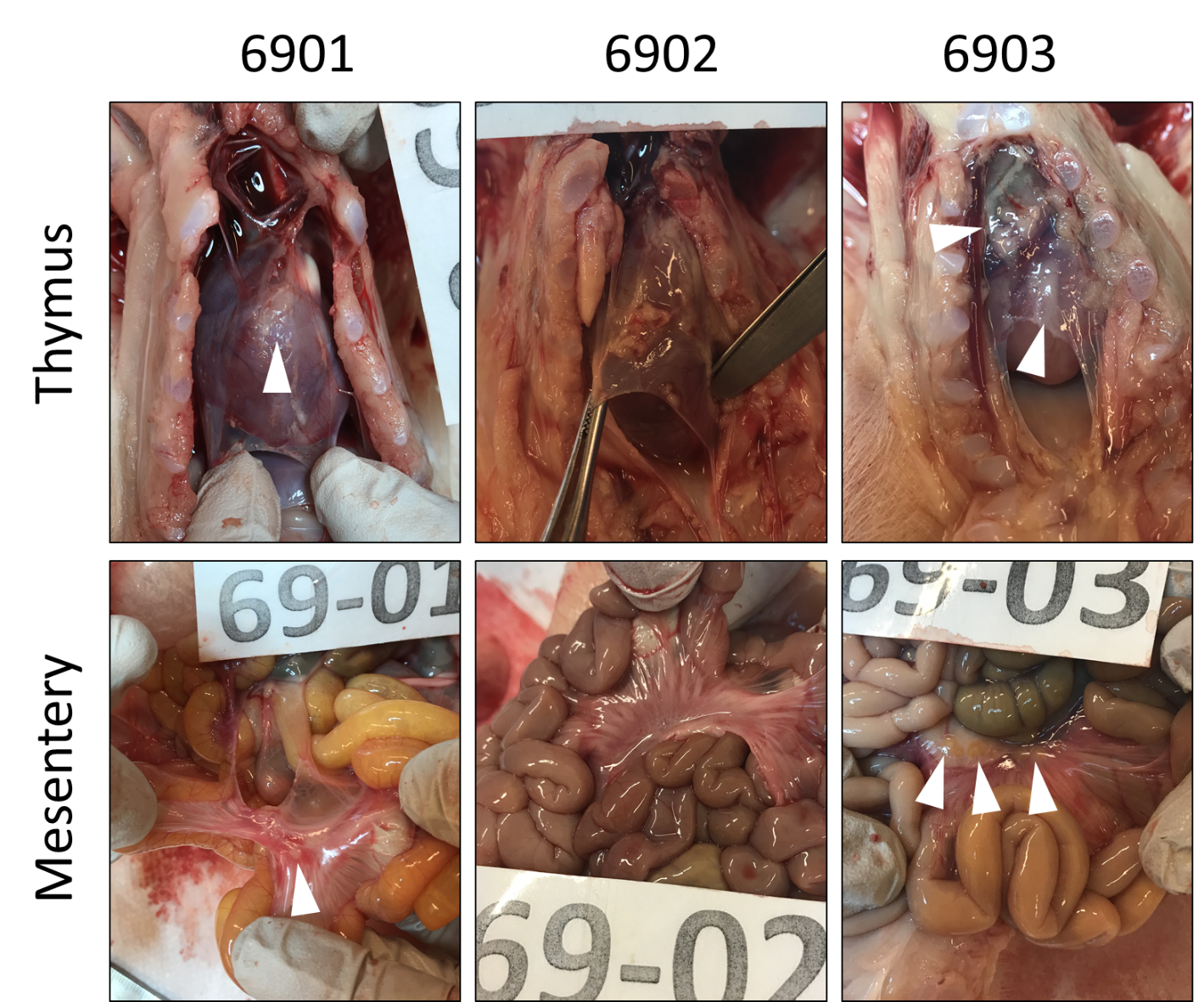

**Supplemental Figure 6. Gross visualization of mesentery and remnant thymic tissue of**

***Art^-/-^ IL2RG^-/Y^* pigs that underwent *in utero* injection of human cells.** Pigs 6901 and 6903 were found to have circulating human leukocytes, while 6902 did not. Some remnant thymic tissue is present on all pigs; 6903 appears to have slightly more. Pigs 6901 and 6903 have lymph node structures present in the mesentery, while 6902 does not. White arrows indicate presence of thymic tissue or mesenteric lymph nodes.

**List of Supplementary Tables**

**Supplemental Table 1A. Artemis genotype and SLA haplotype information for pFF cell lines**

| **Fibroblast ID** | **Sex** | **Genotype** | **SLA Haplotype** | **Additional Information** |
| --- | --- | --- | --- | --- |
| 7709-FB1 | M | 16/+ | 26.6/68.19a |  |
| 7709-FB2 | M | 12/+ | 35.12a/68.19a |  |
| 7709-FB3 | F | 12/+ | 26.6/68.19a |  |
| 7709-FB4 | F | 12/12 | 26.6/35.12a |  |
| 7709-FB5 | F | 12/12 | 26.6/35.12a |  |
| 7709-FB6 | M | 16/+ | 26.6/68.19a | Anticipated SLA donor |
| 7709-FB7 | F | 12/+ | 26.6/68.19a |  |
| 7707-FB1 | M | 12/12 | 26.6/68.19a | Mutagenized line (7707-FB1-U23) |
| 7707-FB2 | M | 12/16 | 26.6/68.19a |  |
| 7707-FB3 | F | 12/+ | 35.12a/68.19a |  |
| 7707-FB4 | M | 12/+ | 35.12a/68.19a |  |
| 7707-FB5 | M | 12/+ | 35.12a/35.12a |  |
| 7707-FB6 | M | 12/+ | 26.6/35.12a |  |

| **Supplemental Table 1B. Definitions of SLA haplotypes** | | | | | | |
| --- | --- | --- | --- | --- | --- | --- |
| **SLA Haplotype** |  |  |  |  |  |  |
|  | **SLA Locus** | | | | | |
|  | **SLA-1** | **SLA-3** | **SLA-2** | **DRB1** | **DQB1** | **DQA** |
| **26.6** | 08:XX | 05:XX | 10:XX | 05:XX | 08:XX | 01:XX |
| **35.12a** | 12:XX,13:XX | 05:XX | 10:XX | 06:XX | 07:XX | 01:XX |
| **68.19a** | 07:XX | 01:XX | 01:XX | 04:XX | 07:XX | 03:XX |

**Supplemental Table 2. The efficacy of CRISPR/Cas9-mediated *IL2RG* mutations in *Art ^-/-^* pFFs**

| **Cell Genotype** | **Sex** | **Mutant alleles** |
| --- | --- | --- |
|  |  | **(Mutated colony/tested colonies)** |
| Art ^-/-^ | Male | 5/202 (2.5%) |

**Supplemental Table 3. Somatic cell nuclear transfer results in embryos for transfer into fertile recipients**

| **Donor cell type** | **No. of embryos transferred** | **No. of ET recipients** | **No. (%)of pregnancies at day 30** | **No. (%) of term pregnancies** | **No. (%) of pigs born** |
| --- | --- | --- | --- | --- | --- |
| Art^-/-^IL2RG^-/Y^ | 920 | 7 | 4 (57) | 2 (30) | 5(0.5%) |
| Art^-/+^ | 512 |  |  |  | 0 |

**Supplemental Table 4. Humanization attempts in single mutant Art^-/-^ pigs by intravenous and intraosseous injection**

| **Animal ID** | **Injection route** | **Cell dosage (cells/kg)** | **Busulfan conditioning** | **Time on trial** |
| --- | --- | --- | --- | --- |
| 202-05 | IV | 217,000 | 2 doses of 2mg Busulfan (2.3 kg) | 10 weeks |
| 202-06 | IV | 271,000 | x | 10 weeks |
| 202-07 | IV | 304,000 | x | 14 weeks |
| 203-01 | IV | 105,000 | x | 10 weeks |
| 203-02 | IV | 105,000 | 2 doses of 4 mg Busulfan (4.7 kg) | 15 weeks |
| 203-05 | IV | 216,000 | x | 15 weeks |
| 202-09 | IV | 111,111 | x | 14 weeks |
| 226-07 | IO | 9,180,000 | x | 7 weeks |
| 227-05 | IV | 17,870,000 | x | 9 weeks |
| 227-07 | IO | 5,900,000 | x | 9 weeks |

**Supplemental Table 5. Somatic Cell nuclear transfer into fertile recipients for *in utero* injection of human cells**

| **Donor cell type** | **No. of embryos transferred** | **No. of ET recipients** | **No. of term pregnancies** | **No. of pigs in utero** | **No. of pigs injected in utero** | **No. of pigs born** | **No. of humanized pigs** |
| --- | --- | --- | --- | --- | --- | --- | --- |
| Art^-/-^ IL2RG^-/Y^ | 320 | 1 | 1 (100%) | 6 | 3 | 3 | 2 |

**Supplemental Table 6. Primers used for genotyping fetal fibroblast lines and cloned piglets**

| **Primer** | **Sequence (5’- 3’)** | **PCR protocol** |
| --- | --- | --- |
| SRY-F | ATGGACGTGAAACTAGAGGAAG | 94˚C for 5 min, 30 cycles of 94˚ for 30s, 60˚C for 30 s, and 72˚C for 35 s, followed by 72˚C for 5 minutes |
| SRY-R | TGTGAGTGACTTAACTGGCTTT |  |
| IL2RG F1 | GGACCAAGAAAGAGGTTAGCCAG | 95˚C for 5 min, 38 cycles of 94˚ for 30s, 58˚C for 30 s, and 72˚C for 30 s, followed by 72˚C for 5 minutes |
| IL2RG R1 | CGGCTTTTGTTTACTTGTCTGTC |  |
| ART16 F | CTCAGTGGGTTAGGGACCTG | 95˚C for 2 min, 38 cycles of 94˚ for 30s, 54˚C for 30 s, and 72˚C for 30 s, followed by 72˚C for 5 minutes |
| ART16 R | GCCATCTGATAGGGTTTCCA |  |
| ART12 F | GCTAAAGTCCAGGCCAGTTG | 95˚C for 2 min, 38 cycles of 94˚ for 30s, 54˚C for 30 s, and 72˚C for 30 s, followed by 72˚C for 5 minutes |
| ART12 R | CAAGAGTCCCCACCAGTCTT |  |

**Supplemental Table 7. Primers used to test potential off-target mutagenesis**

| **Potential target gene** | **Accession** | **Location** | **Target (crRNA, DNA)** | **Primers (F, R) (5'-3')** | **Product length** |
| --- | --- | --- | --- | --- | --- |
| VPS54 | ENSSSCG00000027509 | Chr. 3: 81,716,210-81,779,070 | GGCCACTATCTATTCTCTG-ANGG | TGTGGTCCAACAATCCACAAC | 382 |
|  |  |  | GGCCAaTcTCTATTCTCTGTgTGG | AAAGAGGTTTCCTGAGTGCCAT |  |
| POLH | ENSSSCG00000001683 | Chr. 7: 44,152,193-44,184,675 | GGCCACTATCT--ATTCTCTGANGG | CAAAAACCACAGTCCAGCCC | 261 |
|  |  |  | GGCCACcATCTGGATaCTCTGtTGG | ATAGTCCGAGTGTTCTGGCG |  |
| JPH1 | ENSSSCG00000006174 | Chr. 4: 67,243,406-67,423,310 | G--GCCACTATCTATTCTCTGANGG | CGGGAACCAACGGACGTATAG | 214 |
|  |  |  | GAGGCCACTgTCTgTgCTCTGAAGG | ACTCTCCTCTGTCTTTGGGATAA |  |
| GRIN2B | ENSSSCG00000000614 | Chr. 5: 62,070,371-62,272,186 | GGCCACTAT-CTATTCTCTGANGG | GCTAGTACTTGGTCAGAGATTGGA | 212 |
|  |  |  | GGCCAaTATTaTATTCaCTGATGG | ATGTAGGGTGTGTGTGGTCTG |  |
| VEPH1 | ENSSSCG00000011726 | Chr. 13: 105,238,633-105,486,983 | GGCCACTAT-CTATTCTCTGANGG | GAAGAGTACTAGCTGTTCAAATTGT | 220 |
|  |  |  | GGCtgCTATTCTgTTCTCTGAGGG | ACCTGTTCAGCTTTCCCTGG |  |
| HHLA2 | ENSSSCG00000027275 | Chr. 13: 160,195,017-160,239,674 | GGCCACTATC--TATTCTCTGANGG | GGCAGAAACTGTAGTCTGCTTT | 152 |
|  |  |  | caCCACTATaCTTATTCTCTGATGG | GCTCTTGTCATCGCATACTGTTC |  |
| RASA1 | ENSSSCG00000014146 | Chr. 2: 97,400,410-97,515,957 | GGCCACTATCT--ATTCTCTGANGG | ATTTGGGTGGAGATTGCTCAGT | 214 |
|  |  |  | ttCCACTATCTTAATTCTCaGATGG | TCACAAAGCATCCAGAGAAAGATG |  |
| ASB5 | ENSSSCG00000030395 | Chr. 15: 44,410,073-44,464,546 | GGCCACT--ATCTATTCTCTGANGG | CAGCATCACTTTGTTCCCCA | 279 |
|  |  |  | GGCCACTGCAaCgATTaTCTGAAGG | AGCAGACTCTATGTGGGTAAAGC |  |
| PLPP4 | ENSSSCG00000010691 | Chr. 14: 141,831,969-141,983,156 | GGCCACTATCTAT--TCTCTGANGG | AGAGAGTTGGCATAACACCCTG | 385 |
|  |  |  | GGCCACTATaTAaCCTCTCaGATGG | ACGCTGGCCTTTTCTCTTTTTC |  |
| SLC353 | ENSSSCG00000010162 | Chr. 14: 60,839,431-60,934,051 | GGCCACTATC--TATTCTCTGANGG | CTGAACCAGAACTTCCTACCTCG | 200 |
|  |  |  | aGCCACTAgCTTcATTCTCTGAAGG | GGGAAGGTGGACGAGATGAC |  |
